# Locus-specific proteomics identifies new aspects of the chromatin context involved in V region somatic hypermutation

**DOI:** 10.1101/2022.09.08.507190

**Authors:** GuoJun Yu, Zhi Duan, Yongwei Zhang, Jennifer T Aguilan, Simone Sidoli, Matthew D Scharff

## Abstract

Activation-induced cytidine deaminase (AID) somatically hypermutates the immunoglobulin heavy chain variable region (IGHV) gene to create the antibody diversity required to resist infections. This hypermutational process involves many pathways including transcription, DNA structural change and repair. While many of the proteins involved have been identified, their relative abundance, organization and regulation have not been resolved and additional factors and pathways need to be identified. To identify the proteome occupying IGHV, we have utilized dCas9-APEX targeted by guide RNAs to biotinylate and enrich the proteins associated with the mutating V region chromatin in the Ramos human B cell line and compared them to the non-mutating downstream constant region (C) chromatin. We identified hundreds of proteins specifically enriched on the V or C region. We confirmed the functionality of selected factors by examining the changes in the V region-specific proteome after inhibiting transcriptional elongation and somatic mutation with the Dot1L inhibitor EPZ004777.

**Summary:** Locus-specific proteomics using dCas9-APEX identifies new aspects of the chromatin context involved in V region somatic hypermutation (SHM) in the human Ramos B cell line. An inhibitor of Dot1L which participates in SHM is used to identify functional SHM-related factors.

## Introduction

To survive infections the B cells in our body need to constantly generate antibodies with increasing affinity and changes in fine specificity against pathogens. To accomplish this, the B cells enter into the germinal center where they turn on the expression of activation-induced deaminase (AID) to initiate the somatic hypermutation (SHM) of the variable region (V) genes that encode the antigen binding site. AID deaminates cytosines (dC) in single-stranded DNA (ssDNA) converting them to uracils (dU) that are either replicated to produce C->T mutations or processed by error-prone base excision repair or mismatch repair to produce other sorts of base changes in the V region (Feng et al., 2020; Methot and Di Noia, 2017). While AID is responsible for the mutations in dC on both strands, half of the mutations in the V region are in A:T bases and these arise because the G:U mismatches that are generated by AID can also recruit error-prone mismatch repair to create mutations at A:T (Peled et al., 2008). AID prefers WRC (W = A/T, R = A/G, the underlined nucleotide is mutated) motifs. AID-induced hypermutation is largely restricted to the V region and to switch (S) regions in the Ig locus where it initiates the process of class switch recombination (CSR), although there are some mutations of distant non-Ig “off-target” sites that occur at a much lower mutation frequency (Álvarez-Prado Á et al., 2018; Casellas et al., 2016). B cells expressing V region mutations that increase the affinity of the of antibody are selected for further growth and differentiation resulting in an overall affinity maturation and an increase in the protective efficacy of the antibody response (Victora and Nussenzweig, 2022).

It is widely accepted that the transcription-related processes play a central role in generating ssDNA that is the substrate for AID and in recruiting the factors that are responsible for the mutational process (Bransteitter et al., 2003; Sun et al., 2013). Cis-acting DNA regions, transcription factors, several transcription elongation factors, histone modifications and a histone chaperone which contributes to active transcription have been shown to play important roles in this process (Duan et al., 2021; Feng et al., 2020; Tang and MacCarthy, 2021; Yu et al., 2021a; Yu et al., 2021b). For example, Spt5 was reported to facilitate the transition from pausing to elongation and to promote V region SHM both through its physical interaction with AID and by inducing RNA polymerase II (RNA Pol II) stalling (Methot et al., 2018; Pavri et al., 2010; Pham et al., 2019). Factors involved in processing nascent RNA such as the RNA exosome that regulates transcription elongation by degrading newly transcribed RNA have also been reported to facilitate the recruitment of AID to its targets and to expose ssDNA to AID (Basu et al., 2011). AID mutation of the IgM switch region initiates CSR to change the antibody isotype and many different factors involved in this process including 53BP1, KAP1, PTBP2, FAM72A, and ROD1 have been identified using the proteomics of AID interacted proteins, shRNA screening and genome wide CRISPR/Cas9 screening in the mouse B cell line CH12 (Chen et al., 2018c; Feng et al., 2021; Jeevan-Raj et al., 2011; Manis et al., 2004; Nowak et al., 2011; Willmann et al., 2012). However, except for AID itself, the individual depletion of these and other physically or functionally associated factors do not fully eliminate SHM on either V or S regions and all of the proteins identified so far also have genome wide functions. Furthermore, in spite of these and many other studies, we still do not have a clear and complete model of how AID mutational activity is highly restricted to the Ig V and S regions of the Ig gene.

The Ramos germinal center like Burkitt’s lymphoma cell line has proven to be a good system to study the role of individual candidate genes and processes such as transcription as well as the role of larger genomic regions such as topologically associating domains (TADs) in the SHM of the human heavy chain V region (Dinesh et al., 2020; Duan et al., 2021; Senigl et al., 2019; Tarsalainen et al., 2022; Yu et al., 2021b). In order to obtain a broader more quantitative and less biased view of chromatin environment in which AID induced mutation is occurring in situ, we used Mass Spectrometry that would allow us to compare and quantify both known and unknown proteins that were associated with the chromatin in the mutating V and non-mutating C regions. With the advancement of Mass Spectrometry with high resolution, several seminal methods had been developed to study the protein context at a specific genomic locus (Gauchier et al., 2020). In one such system, dead Cas9 has been conjugated with the ascorbate peroxidase APEX (dCas9-APEX) (Gao et al., 2018; Myers et al., 2018). The dCas9-APEX protein is delivered through the use of a guide RNA (gRNA), and then the APEX enzyme can be activated to label the proteins located on this specific chromatin region with biotin in live cells. The biotinylated proteins can then be pulled down and analyzed by Mass Spectrometry to identify the proteome in the genomic locus. This technique has been largely used to study the protein complexes on repeated genomic regions like centromeres or telomeres and at single gene loci, but as far as we can tell it has not been used to distinguish the protein environment on chromatin on the two regions that are part the same gene, like the V and C region of the Ig locus. Moreover, this technique has not been used in combination with transcription inhibitors treatment to dissect the dynamic changes in chromatin context in cells.

In the current study, in order to identify a broad spectrum of proteins that mainly play roles in the V region that is mutated by AID rather than the downstream C region that does not undergo AID-induced mutation, the dCas9-APEX platform with specific gRNAs that target the human heavy chain V and the C region, separately, was established in the previously described Ramos Rep161 cell line (Wang et al., 2014). Using the dCas9-APEX with Mass Spectrometry, we are able to distinguish the different protein contexts on V compared to the C regions that are only 8 kb apart. We identified, as reported previously (Feng et al., 2020), proteins involved in transcription elongation and RNA splicing to be enriched on V region, as well as additional factors involved in these processes. Furthermore, we could compare changes in protein relative abundance between the V and C region when the Dot1L inhibitor was added to perturb elongation and mutation, confirming the potential roles of transcription elongation and RNA splicing pathways in SHM of V region. Finally, based on Mass Spectrometry data, we found that the super elongation complex also plays a synergistic role with Dot1L pathway in regulating SHM of V region in Rep161 cell line.

## Results

### Successful establishment of the dCas9-APEX platform in Rep161 Ramos cell line

In order to identify and quantify the specific protein complexes on the V region using the dCas9-APEX system, as shown in Fig. 1A, we stably transfected Rep161 cell line with a vector expressing doxycycline driven Flag-tagged dCas9 conjugated to C terminus to the APEX enzyme linked by a T2A peptide to a EGFP protein (Myers et al., 2018). T2A is cleaved after translation to release the EGFP. Due to the low transfection efficiency of B cell lines, we single-cloned the stably transfected cells in 30 96-well plates and selected all of the clones that expressed the EGFP under doxycycline induction and created a pooled culture that was used in all of the experiments described in this paper. Single guide RNAs (gRNA) that individually targeted the variable or constant regions of the Ramos REP161 heavy chain locus were designed. In the Rep161 cell line, the mCherry sequence is inserted in frame at the 5’ end of the endogenous IGHV gene in Rep161 (see cartoon below Figure 1C and D and in Figure 2I) so as to use the loss of mCherry to quantify mutation (Wang et al., 2014). Those studies had shown that the mCherry and the V region undergo AID mutation at approximately the same frequency. In order to largely avoid the targeting of non-rearranged V regions, the gRNA for V region was designed to target the “unique” 3’ end of mCherry where it is fused to the endogenous V region. Since others have estimated that the biotin reaction has an ~ 400bp resolution flanking both sides of the gRNA targeted site (Myers et al., 2018), this should allow us to biotinylate much of the fused mCherry-V region (1.2 kb). The gRNA for Cμ region targets the exon 4 of the constant μ gene which is approximately 8 kb downstream from the V region gRNA targeted site Figure 1A, second panel). In order to achieve high transfection efficiency of gRNAs, we used the third generation lenti virus vector for gRNAs and separately stably infected the dCas9-APEX expressing cells with the virus expressing each gRNA. This provided us with two different cell lines which are expressing either dCas9-APEX plus V region gRNA (named “V-gRNA”) or dCas9-APEX plus C region gRNA (named the “C-gRNA” cell line) (Fig. 1A, “Transfection” panel).

**Figure 1.**
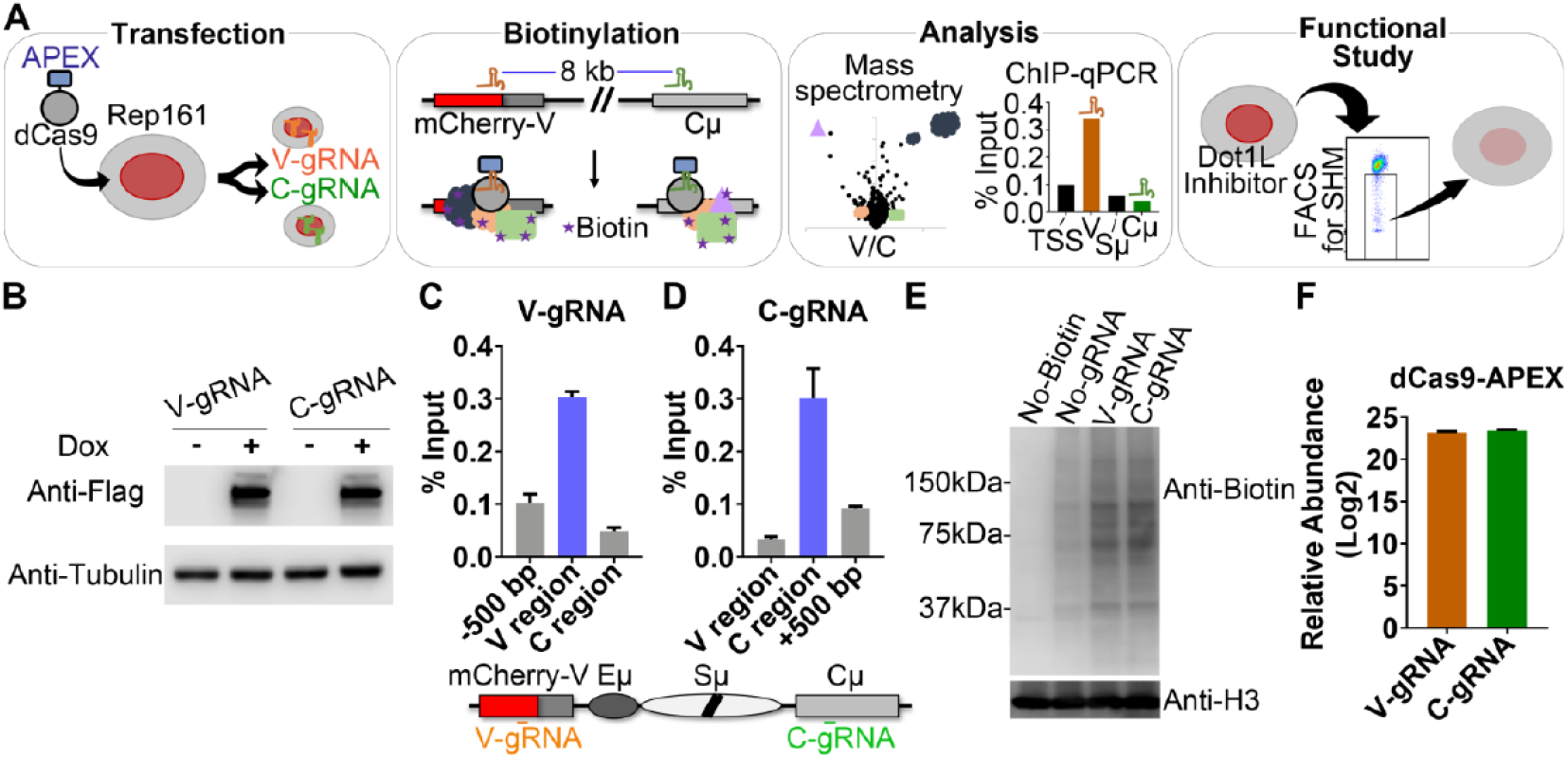
Successful establishment of dCas9-APEX platform in Rep161 cell line. (A) Schematic pipeline of the study. (B) Western blotting of Flag-tagged dCas9-APEX that is stably transfected in Rep161 cells. (C-D) ChIP-qPCR of Flag-tagged dCas9-APEX using anti-Flag antibody for V-gRNA and C-gRNA stably transfected cells, respectively. In the ChIP experiments throughout the paper, the results were normalized to the percentage of input DNA and an IgG negative control was subtracted. Each ChIP was done in triplicate and the error bars represent Standard Error of the Mean (SEM) of those determinations. (E) Western blotting of biotinylated proteins of purified chromatin fraction from each group (see Methods). (F) The fold enrichment of dCas9-APEX protein on total chromatin in both V-gRNA and C-gRNA transfected cells compared with the cells with only dCas9-APEX transfection.

**Figure 2.**
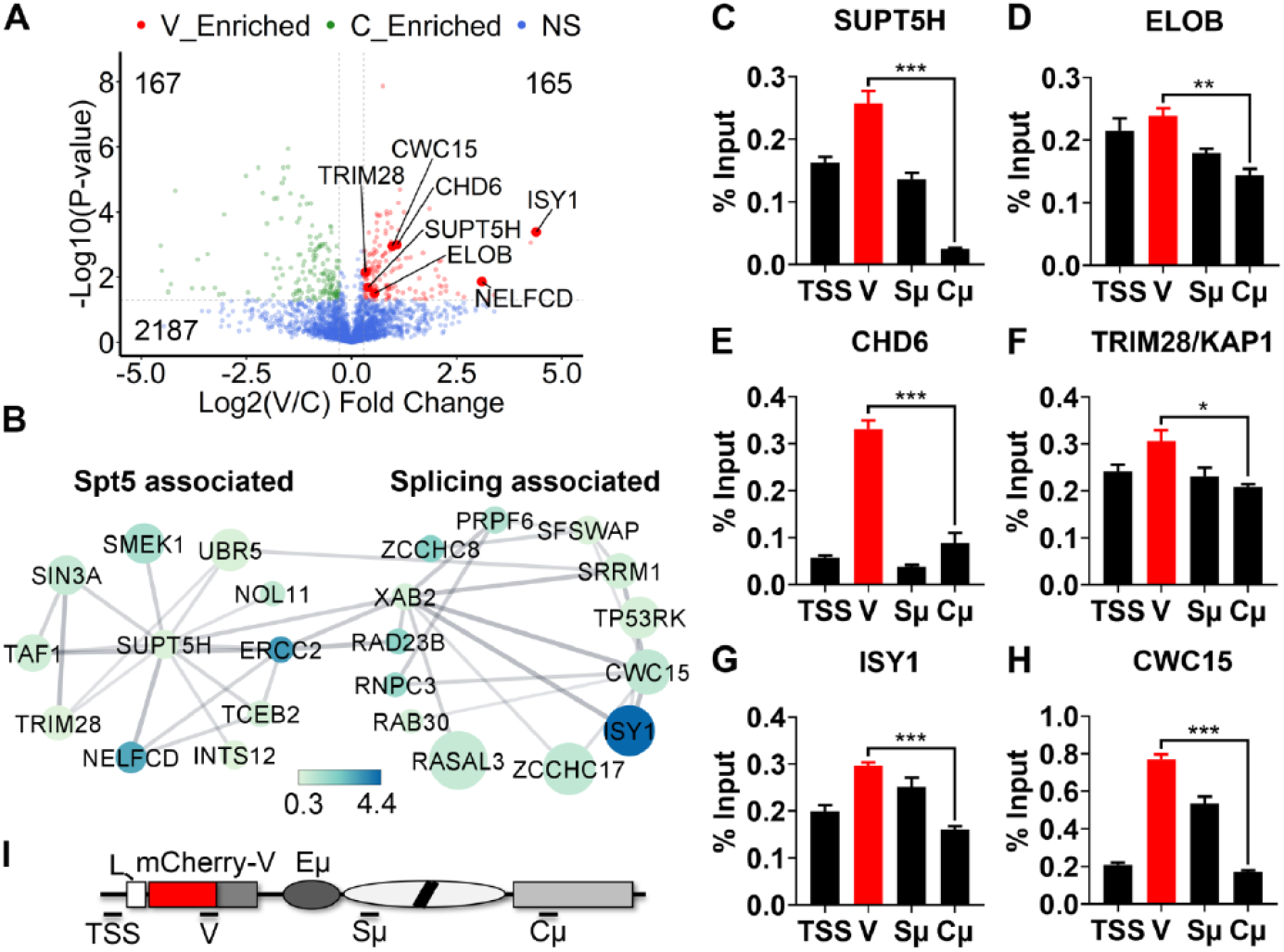
Identification, functional interactome and ChIP verification of proteins that based on APEX-Mass Spectrometry are enriched on V region compared with the C region. (A) Volcano plot showing the fold change comparison between V and C regions on the X axis and the p-value on the Y axis. Representative proteins enriched on the V region involved in transcription and splicing were labeled with their respective names. Three independent Mass Spectrometry analyses were performed for each group. (B) Functional network analysis by STRING of Spt5 and RNA splicing related proteins that are statistically significantly enriched on V region compared to the C region. The size of the circle represents the P-value with bigger circles reflecting greater significance. The color indicated by the color band below the interactomes represents the fold difference in abundance of each protein on the V region compared with the C region. (C-H) ChIP-qPCR verification of representative proteins that are more abundant on the V region compared to the C region. Each ChIP was done in triplicate and the error bars represent Standard Error of the Mean (SEM) of those determinations. (I) A schematic of the mCherry-V region construct used to replace the endogenous IGHV4-34.01 gene in the Rep161 Ramos reporter cell line showing the location of sites used for ChIP-qPCR in panels C-H. In the current and following figures, * means p-value <= 0.05, ** means p-value <= 0.01, *** means p-value <= 0.001.

Based on western blotting (WB) with anti-Flag, the expression level of dCas9-APEX in V-gRNA and C-gRNA cell lines is similar after 60 hours of doxycycline induction (Fig. 1B). We treated the cells for 60 hours for all the experiments, because we barely observed the EGFP signal after 24 hours of doxycycline treatment in the dCas9-APEX selected clones. Then, in order to confirm that the dCas9-APEX protein had been delivered to the targeted site by each gRNA, we treated each cell line with doxycycline and performed ChIP-qPCR with anti-Flag antibody. Fig. 1C shows that the Flag-tagged dCas9-APEX was delivered to the mCherry-V region by gRNA targeting the 3’ end of the mCherry compared to a region 500 bp upstream of the gRNA targeted site and to the C region which is around 8 kb downstream. The dCas9-APEX was guided to the constant region by the C region gRNA targeting exon 4 of Cμ compared with the 8kb upstream V region and a region ~500bp downstream from the gRNA targeted site (Fig. 1D). The relatively weak signal observed in the region 500 bp upstream of the mCherry gRNA targeted site and 500 bp downstream of the Cμ gRNA targeted site is consistent with the size of the sonicated fragment for ChIP which is normally between 300bp and 1 kb. The above data clearly demonstrated that the dCas9-APEX can be efficiently and selectively delivered to the respective gRNA targeted site and the delivery efficiency of dCas9-APEX to each region was comparable.

Since these studies suggested we could achieve relatively specific biotinylation of the proteins associated with the V and C region chromatin, separately, we then performed the biotinylation experiment in situ (Fig. 1A, “Biotinylation” section) for each live cell line including the cells only expressing the dCas9-APEX (Myers et al., 2018). The relevant published studies normally used extracts of whole nuclei for the pulldown and Mass Spectrometry determination. However, since this could introduce nonspecific signals due to the non-binding dCas9-APEX located in the nuclei or even perhaps labeling of a protein such as AID which is in huge excess in the cytoplasm (Feng et al., 2020), we extracted the native (non-cross linked) chromatin fraction by removing the cytoplasmic and nucleoplasm protein fraction so that only the proteins that bound to the chromatin would be collected for the pull-down. As shown in the WB with the anti-biotin antibody (Fig. 1E), the total native chromatin fraction from V-gRNA and C-gRNA cell lines contain more detectable biotinylated proteins compared with the dCas9-APEX cells without any gRNA and the dCas9-APEX cells without a biotinylating reaction. This suggests that the dCas9-APEX can significantly modify proteins on chromatin under the direction of specific gRNA.

This was confirmed when we performed the pull-down with streptavidin-conjugated magnetic beads of the native chromatin fraction from V-gRNA and C-gRNA cell lines after the biotinylation reaction. The pull-down from these two cell lines and their respective input chromatin faction were subjected to Mass Spectrometry identification with independent biological triplicates for each group. From the proteomic data, we found the relative abundance of the dCas9-APEX protein on chromatin from the input of each cell line was similar (Fig. 1F) which further confirmed the data shown in Fig. 1B-D and clearly demonstrated that the dCas9-APEX can be delivered to each locus by the corresponding gRNA comparably. Overall, the above data proves that the dCas9-APEX system had been successfully established in our Rep161 cell line.

### Quantitative analysis of identified proteins that are enriched on V region compared with the C region

To capture Mass Spectrometry data from both cell lines (V-gRNA and C-gRNA), we normalized the pull-down data for each protein with its respective input and then compared the data from V-gRNA cell line with the C-gRNA cell line (see Methods). When the Log2(V/C_Fold_Change) >= 0.3 and the p-value <= 0.05 (-Log2(p_value) >= 4.3) we classified the protein as significantly enriched on V and when Log2(V/C_Fold_CHange) <= −0.3 and the p-value <= 0.05 (-Log2(p_value) >= 4.3) the protein was classified as enriched on the C region. Based on these criteria, 165 proteins are significantly enriched on the V region, 167 proteins are significantly enriched on the C region and 2,187 proteins were associated with the Ig locus chromatin but did not differ between V and C region (Fig. 2A).

Several factors had been reported to play roles in regulating SHM of the IGHV gene but their relative abundance on the V region remains unclear either because there are no antibodies that can be used for ChIP or the differences in the affinity of those antibodies that do work makes it impossible to compare the abundance of different proteins. In ChIP studies the chromatin is usually cross-linked. However, we found that the backgrounds of the Mass Spectrometry analysis were much higher if we used cross-linked rather than native chromatin, so we have not cross linked the chromatin in the studies described here. This may result in some proteins being lost because their binding to chromatin is relatively weak (Gauchier et al., 2020).

Totally, we identified 2,519 proteins associated with V and C regions including AID and 10 other proteins that are known to play roles in SHM (Table S1). Beta-catenin-like protein 1 (CTNNBL1) (Ganesh et al., 2011), polypyrimidine tract-binding protein 2 (PTBP2) (Nowak et al., 2011) and polypyrimidine tract-binding protein 3 (PTBP3/ROD1) (Chen et al., 2018c) all interact with AID to regulate SHM. The FACT complex (SPT16 and SSRP1) (Aida et al., 2013), germinal-center associated nuclear protein (MCM3AP) (Singh et al., 2013), HIRA and transcription elongation factors (SPT4, SPT5 and SPT6) regulate SHM of region by modulating the transcription (Wang et al., 2014; Yu et al., 2021b). Based on our Mass Spectrometry data, none of these are significantly more abundant on the V region than the C region except Spt5 and possibly HIRA (Table S1). In particular AID was equally abundant on the V and C region chromatin even though the C region does not undergo detectable AID-induced mutation. It is important to note that this assay does not reveal whether the AID associated with the C region is making direct contact with the DNA. Nevertheless, it is surprising and provides new important information on the complexity of the regulation of V region hypermutation and demonstrates the value of documenting the relative abundance using dCas9-APEX-Mass Spectrometry to study the occupancy of interesting factors on a specific genomic locus.

In Fig. 2A, the volcano plot shows the distribution of the Log2(V/C_Fold_Change) of proteins that differ significantly in their abundance at the V and C with a p-value (-log10 form). The red dots show the 165 proteins that are enriched on the V region while the green dots show the 167 proteins that are enriched on the C region. Some of the proteins enriched on V region are related to the transcription initiation and elongation and RNA splicing and are named in Fig. 2A. One important example is NELFCD which plays a critical role in the transition from transcriptional pausing to elongation (Chen et al., 2018b) but has not been identified in previous studies as playing a role in SHM. Based on functional network analysis using the STRING program (Szklarczyk et al., 2021) in Cytoscape (Shannon et al., 2003), there are many additional proteins that are enriched on the V region that are associated with the Spt5 (SUPT5H) which is a transcription elongation factor known to play a role in SHM (Fig. 2B). There are also many proteins enriched on the V region that like ISY1, that has not been reported before, that are associated with RNA splicing (Fig. 2B) confirming that these two processes could play important roles in V region mutation. These findings indicate that dCas9-APEX-Mass Spectrometry can reveal additional proteins in pathways known to be involved in SHM and potential interactions between complexes that contribute to the somatic hypermutation on V region.

In order to independently confirm the relative abundance of the proteins that appear to be enriched by the dCas9-APEX technique on the V region compared to the C region, we searched for antibodies that could be used for ChIP-qPCR. In the end, we were able to perform reliable ChIP-qPCR for 6 apparently enriched targets on V region (Fig. 2C-H). The transcription start site (TSS) which is ~900 bp upstream of region (Wang et al., 2014) and the switch μ (Sμ) region that is mutated by AID in Ramos (Qian et al., 2014) were included in the ChIP-qPCR and all 6 of these candidates are significantly enriched on V region compared with the C region. Spt5 and ELOB are involved in the transcription elongation (Chen et al., 2018b; Garrett et al., 1995). CHD6 is a chromatin remodeler and regulates transcription (Manning and Yusufzai, 2017). KAP1 is a histone methyltransferase for the K9 residue on histone H3 and plays role in class switch recombination partially by interacting with AID (Jeevan-Raj et al., 2011). ISY1 and CWC15 are RNA splicing factors and CWC15 is reported to interact with CTNNBL1 (van Maldegem et al., 2015). Although generally the ChIP-qPCR data for these proteins is consistent with the Mass Spectrometry data, the fold enrichment between V and C by ChIP-qPCR is different from Mass Spectrometry data. For example, the fold change of ISY1 between V vs C is ~8 fold by Mass Spectrometry but only 2 fold by ChIP-qPCR. This discrepancy could be due to the crosslinking used in the ChIP but not in the dCas9-APEX-Mass Spectrometry, the size of the fragments used for ChIP and/or accessibility to antibody for ChIP-qPCR. Except for the Spt5, which is known to regulate SHM of V region through its role in elongation (Wang et al., 2014), the exact roles of the other 5 factors in SHM remain unclear. The fact that the ChIP-qPCR data was in a general sense consistent with the Mass Spectrometry data suggests that the dCas9-APEX is a useful reporter of the relative abundance of proteins in our Rep161 cell line. This was important because previous reports mainly used this technique to determine the protein context in regions like the telomeres and centromeres which are large and contain many sequence repeats, while in the current study we successfully adapted this technique to distinguish the protein components on the two relatively close regions on the same gene locus which are only 8 kb apart.

### Quantitative analysis of proteins that are enriched on C region compared with the V region

167 proteins are enriched on the C region that does not undergo detectable SHM and these proteins were shown in green dots in Fig. 3A which is the same volcano plot shown Fig. 2A except that representative proteins enriched in the C region are labeled with their respective name in Fig. 3A. DNMT3A is an enzyme responsible to the DNA methylation and is significantly enriched on the C region compared with the V region suggesting that the DNA sequence in C region may be DNA methylated even though C region is highly transcribed. Previous studies reported that the constant region sequence of an Ig locus is methylated which is consistent with our Mass Spectrometry data for DNMT3A (Oudinet et al., 2019) and for a number of proteins which based on functional network analysis using the STRING program in Cytoscape are associated with DNMT3A (27 proteins, Fig. 3B). Similarly, there were 11 proteins enriched in the C region which based on the STRING program are associated with BRCA1 related proteins (BRAP, BRAT1 and BARD1, Figure 3B). Histone macro-H2A.1 (H2AFY) and the Histone PARylation factor 1 (HPF1) are also enriched on C region and these proteins are involved in many functions including DNA repair (Chen et al., 2018a; Sun and Bernstein, 2019; Suskiewicz et al., 2020). HMGB2 is particularly interesting because it facilitates binding of DNA to increase interactions between cis-acting proteins for example in the repair of double-stranded DNA breaks (Voong et al., 2021). The enrichment of DNA repair proteins on the C region raises the possibility that the C region may be mutated by AID but the mutations are repaired with high fidelity by DNA repair systems. As already noted, in the Rep161 cells, based on Mass Spectrometry the abundance of AID associated with the V and C regions is roughly the same (Table S1), so this becomes a real possibility that needs to be studied in the future.

**Figure 3.**
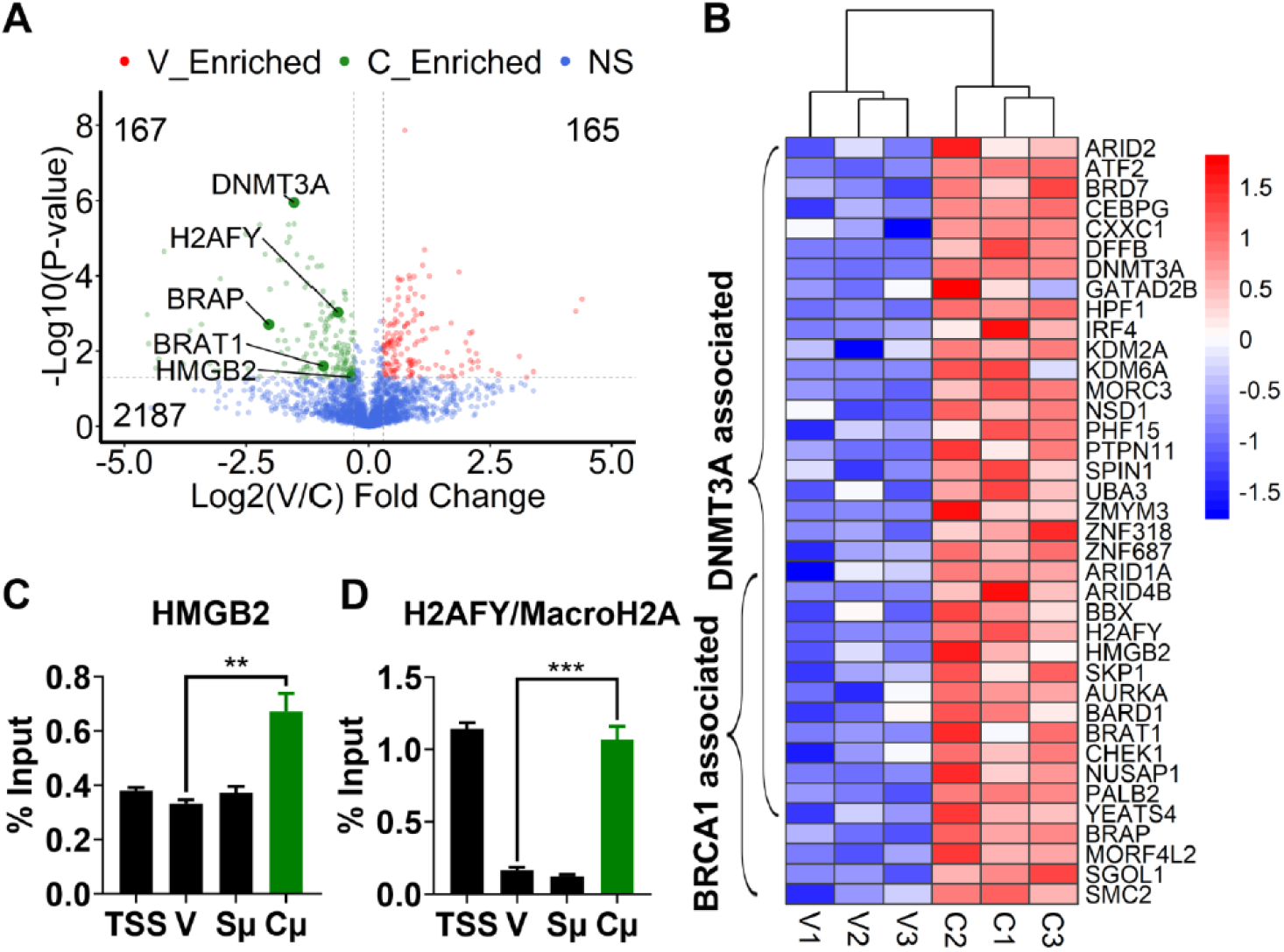
Identification, functional interactome and ChIP verification of proteins that based on APEX-Mass Spectrometry are enriched on C region compared with the V region. (A) The volcano plot showing the fold change comparison between V and C regions with p-value distribution is the same plot shown in Figure 2 A except that here representative enriched proteins on the C region were labeled with their respective names. Three independent Mass Spectrometry analyses were performed for each group. (B) A heat map reflecting the functional network analysis by STRING of DNMT3A and BRCA1 associated proteins that are enriched on the C region. The vertical bar on the right shows relative enrichment in the form of Z-score. (C-D) ChIP-qPCR verification of the representative proteins that are abundant on C region compared with V region. Each ChIP was done in triplicate and the error bars represent Standard Error of the Mean (SEM) of those determinations. The mCherry-V region construct and the locations of the sites used for ChIP-qPCR are the same as in the cartoon in Figure 2.

We found only two antibodies that allowed reliable ChIPs of the proteins that were enriched in the C region compared to the V region and these were against high mobility group protein B2 (HMGB2) and H2AFY/MacroH2A. As shown in Fig. 3C-D, by ChIP-qPCR the HMGB2 and H2AFY/MacroH2A are more abundant on the C region than the V region which is consistent with the Mass Spectrometry data. Interestingly, the abundance of H2AFY on the TSS which is ~900 bp upstream of V region is comparable with its abundance on the C region. In addition to its other functions noted above, H2AFY can either promote or repress the active transcription in different situations. We have previously found that the RNA polymerase II is more abundant on the TSS of Ig locus in Rep161 than the C region (Duan et al., 2021; Yu et al., 2021b) so it is possible that H2AFY may have different roles on the V and C regions.

To further confirm the Mass Spectrometry data, we performed ChIP-qPCR of selected proteins that had the same abundance on the V and C regions as shown in Fig. S1. By Mass Spectrometry, histone-binding protein RBBP4, RuvB-like 2 and DNA replication licensing factors (MCM3 and MCM5) have similar abundance between V and C regions and this was confirmed by ChIP-qPCR.

In summary, the above data clearly demonstrated the ability of dCas9-APEX to reveal differences in the abundance of many proteins in two nearby genomic loci in the human heavy chain locus. This method greatly increases the identification of factors that might participate in the restriction of AID-induced mutations. Most importantly, these studies raised a number of new questions including whether the C region is actually mutated by AID but repaired with high fidelity.

### The protein context on V region determined by dCas9-APEX changed in Rep161 after Dot1L inhibitor treatment

The Mass Spectrometry of the V and S regions provides a snapshot of the presence and relative abundance of a large number of proteins on the chromatin of those regions, but we know that processes that are occurring like transcriptional pausing and elongation are dynamic while many of the proteins we have identified by Mass Spectrometry may play more static though important structural roles. In order to identify some of the more dynamic factors, we used dCas9-APEX to compare the chromatin environment in the presence and absence of EPZ004777, a specific inhibitor of Dot1L-mediated transcriptional elongation (Deshpande et al., 2014). We previously showed (Duan et al., 2021) that Dot1L plays a significant role in SHM but dCas9-APEX provided an opportunity to document the broader dynamic changes in the chromatin environment in situ at the V and C region after inhibitor treatment. To examine this, V-gRNA and C-gRNA cells that had been treated with 4-OHT to induce AID transport into the nucleus for 7 days and with doxycycline (0.1 μg/ml) for the last 3 days of 4-OHT treatment to induce dCas9-APEX were also treated with 20 μM EPZ004777 (dissolved in DMSO) or only DMSO for the whole 7 day period, separately (Fig. 4A). Under these conditions, SHM in the V-gRNA cell line was significantly decreased as determined by flow cytometry (Fig. S2A-B), which is consistent with our previous study (Duan et al., 2021). As noted in our previous study, based on ChIP-qPCR this decrease in mutation was associated with a decrease in the abundance of H3K79me2/3 and an increase in H3K79me1 and a small but statistically significant decrease in total H3 (Fig. S2C-E) on the V region.

**Figure 4.**
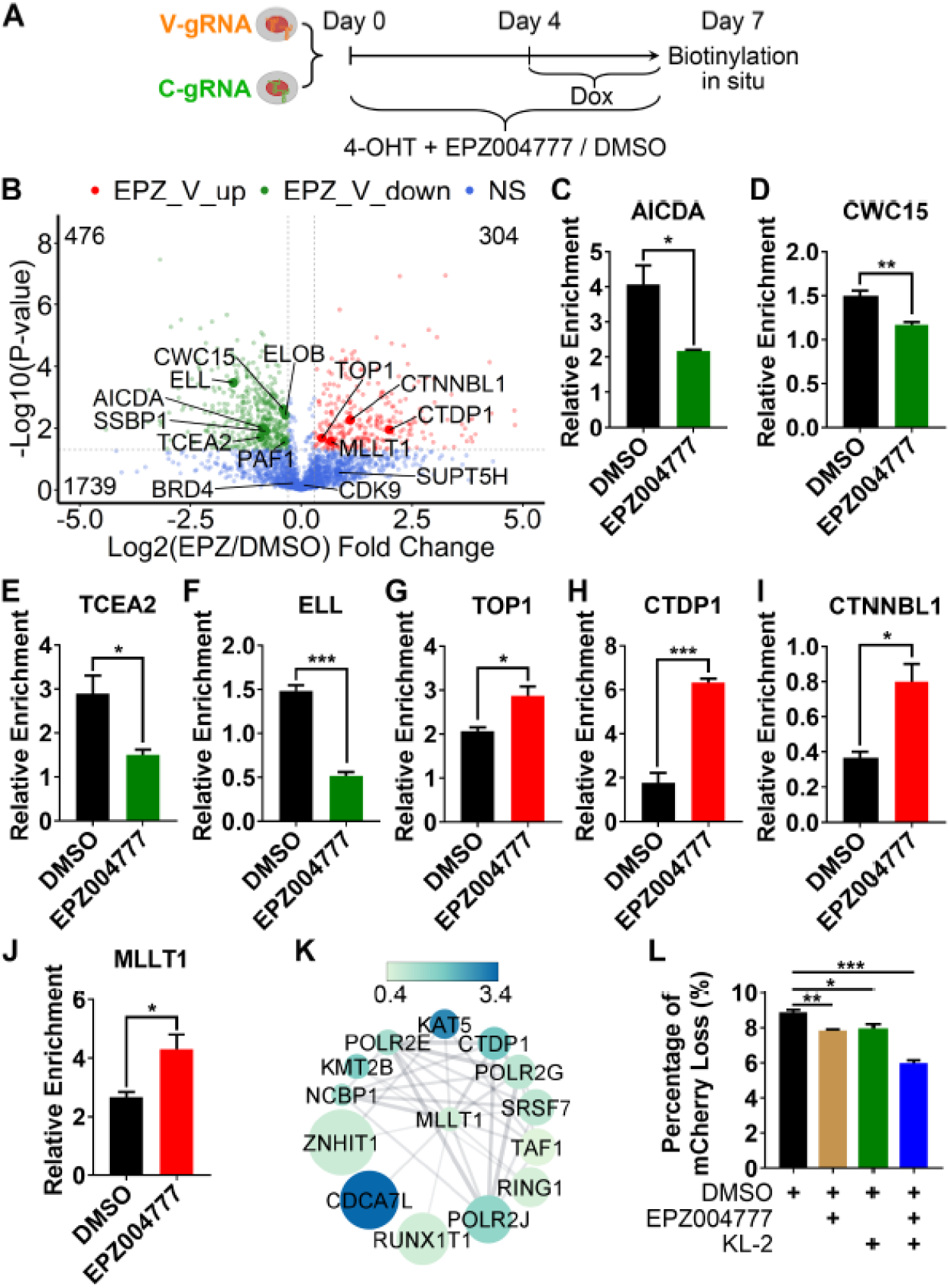
The change of protein abundance on the V region in Rep161 treated with Dot1L inhibitor that is determined by dCas9-APEX. (A) Schema displaying the V-gRNA and C-gRNA cell lines treated with various molecules for Mass Spectrometric identification. (B) Volcano plot showing the fold change comparison of proteins on V region between Dot1L inhibitor EPZ004777 (EPZ) treatment and DMSO control with p-value distribution. Selected proteins that were decreased on the V region by EPZ treatment were labeled with their respective names and are shown in the left-hand side of the volcano plot. These proteins are mainly related to transcription elongation and splicing. Selected proteins that were increased on the V region by EPZ treatment are shown on the right-hand side of the volcano plot and were labeled with their respective names and are mainly related to transcription. (C-F) Bar graphs of representative proteins with decreased relative abundance on the V region based on Mass Spectrometry after EPZ004777 treatment. Error bars represent Standard Error of the Mean (SEM) of three independent Mass Spectrometry determinations. (K) Proteins that are associated with MLLT1 are significantly enriched on the V region after treatment with the Dot1L inhibitor EPZ004777. 13 proteins were identified by STRING to associate with MLLT1. The color bar shows the Log2 fold change of these proteins on the V region after EPZ004777 treated V-gRNA versus the DMSO control. The size of the ball reflects the –Log10 form of P-value. The bigger the ball, the more significant the enrichment on V region after EPZ004777 treatment. (L) The effect of super transcription elongation inhibitor KL-2 on SHM. Both Dot1L inhibitor EPZ004777 (5 μM) and super elongation complex inhibitor KL-2 (0.5 μM) can slightly but significantly decrease the SHM measured by FACS. The error bar represents the SD of three independent experiments. The combination of EPZ004777 (5 μM) and KL-2 (0.5 μM) can decrease the SHM to a greater extent than the individual treatments.

In the C-gRNA cell line, the Dot1L inhibitor treatment decreased the abundance of both H3K79me2/3 and H3K79me1, and again there was a small but in this case not significant decrease in total H3 (Fig. S2F-H). All of these data are consistent with the previous study of the Rep161 cells that did not contain the dCas9-APEX or gRNA and clearly indicate that the Dot1L inhibitor changed the chromatin context of both the V and C regions separately in the V-gRNA and C-gRNA cell lines in ways that were similar to what it did in cells that did not contain dCas9-APEX and gRNA. This suggested that inhibitors could be used with dCas9-APEX to carry out functional chromatin studies, although to the best of our knowledge this had not been previously done.

To examine whether other factors were affected by the Dot1L inhibitor, V-gRNA and C-gRNA cells were collected, biotinylated and the proteins pulled down by streptavidin beads were identified by Mass Spectrometry as in the previous experiments. The previous study showed that the Dot1L inhibitor treatment did not affect the abundance of the BRD4 and CDK9 transcription factors on the Ig locus (Duan et al., 2021) and this is confirmed by Mass Spectrometry (Fig. 4B). EPZ004777 treatment did not affect the abundance of dCas9-APEX protein at the V region compared with the DMSO control (Fig. S3A). In total, 2,519 proteins were identified in the chromatin of V-gRNA cell line including both EPZ004777 (dissolved in DMSO) and DMSO only treated groups. In Fig. 4B, the volcano plot shows the distribution of proteins with Log2 fold change and their −Log10 p-value when comparing Dot1L inhibitor treated group of proteins with the DMSO only group. The green dots represent those 476 proteins whose abundance was decreased on V region after the EPZ004777 treatment, while the red dots represent the 304 proteins that are increased on the V region. In Fig. 4B (left side), we highlighted some downregulated proteins on V region that are involved in transcription elongation and splicing, while on the right side of Fig. 4B we show some representative upregulated proteins on the V region involved in histone methyltransferase, transcription initiation and super elongation. Based on the Mass Spectrometry results now illustrated as bar graphs, after Dot1L inhibitor treatment, the abundance of AID, splicing factor CWC15, and elongation factors TCE2 and ELL were all decreased on the V region (Fig. 4C-F), suggesting that the inhibitor decreased SHM on V region in part by significantly decreasing the abundance of AID itself on the V region and also by decreasing factors involved in transcription elongation. In our previous work (Duan et al., 2021; Yu et al., 2021b) we were unable to perform ChIP to check the AID level on V region due to the lack of an antibody that could reproducibly ChIP for AID in Ramos cells, but we can see it here by Mass Spectrometry (Fig. 4C and Table S1). This illustrates how Mass Spectrometry makes it possible to identify critical proteins that are present transiently or in small amounts or for which there are no realiable antibodies for ChIP.

Based on the Mass Spectromy, DNA topoisomerase 1 (TOP1) was increased on V region after the EPZ004777 treatment (Fig. 4B and 4G). One study had shown that the level of single-stranded DNA (ssDNA) positively correlated with the AID-induced mutation and TOP1 decreased the level of ssDNA (Kobayashi et al., 2009; Parsa et al., 2012). The findings here confirm that increased TOP1 is associated with the decrease in elongation and could result in decreased ssDNA on V region. The RNA polymerase II subunit A C-terminal domain phosphatase (CTDP1, Fig. 4H) was also increased on V region and it is involved in transcription initiation (Friedl et al., 2003). CTNNBL1 interacts with AID to regulate SHM and the MLLT1 component of super elongation complex (SEC); we found CTNNBL1 to be increased on the V region after EPZ004777 treatment (Fig. 4I and J) (Luo et al., 2012). These factors are all involved in active transcription and could represent reciprocal control of initiation or compensation for the inhibitory effect on transcription elongation by the Dot1L inhibitor. Functional interaction network analysis by STRING (Szklarczyk et al., 2021) revealed that 13 proteins associated with MLLT1 that are upregulated on V region after EPZ004777 treatment are (Fig. 4K), suggesting these proteins may form a functional complex to antagonize or compensate for the effect of Dot1L inhibitor and provide new targets for further investigation.

Previous studies suggest that the Dot1L may facilitate active transcription through two different pathways (Cao et al., 2020; Duan et al., 2021). One depends on its enzymatic activity for H3K79 methylation and the other relies on its scaffolding function that allows it to participate in the SEC (Cao et al., 2020). Our recent paper clearly shows that the EPZ004777 treatment did not affect the abundance of Dot1L on the V region suggesting the inhibitor treatment only affects its enzymatic activity but not its scaffolding function (Duan et al., 2021).

In the current study, we observed that 13 proteins associated with MLLT1 (Fig. 4K) were significantly increased on the V region after inhibitor treatment, suggesting the SEC may also play a role in SHM. To test this, we treated the Rep161 cells with 4-OHT to induce mutation and either a low dose of EPZ004777 (5 μM) or of KL-2 (0.5 μM) which is an inhibitor of the SEC (Liang et al., 2018), and with EPZ004777 (5 μM) plus KL-2 (0.5 μM) for 7 days. As shown in Fig. S3C and Fig. 4L, the low dose treatment with either EPZ004777 (11.6% decrease) or the KL-2 (10.2% decrease), that were used to avoid toxicity of both drugs, significantly decreased the SHM. As expected, the individual decreases in mutation were minor, but do suggest that SEC does play a role in regulating the SHM. However, the combination of EPZ004777 and KL-2 which can inhibit both the Dot1L enzymatic activity and the SEC activity gave at least an additive effect significantly decreasing SHM by 32.5% suggesting that they may be functioning as independent pathways (Fig. S3C and Fig. 4L).

### Functional annotation of significantly changed proteins on the V region by Dot1L inhibitor treatment

The proteins that were significantly increased and decreased on the V region after EPZ004777 inhibitor treatment were separately subjected to the “Molecular Function” annotations in Gene Ontology (Ashburner et al., 2000). Fig. 5A shows the top 15 enriched “Molecular Function” annotations in the proteins that are increased on V region after Dot1L inhibitor treatment, while Fig. 5B shows the top 15 enriched annotations from the proteins that are decreased on V region after EPZ004777 treatment. Many of the most highly represented pathways are found in both panels A and B and probably relate to basic chromatin structure. EPZ004777 significantly decreased the abundance of 45 proteins on V region involved in “Transcription factor binding”, “Transcription coregulator activity” and “Transcription coactivator activity” (Fig. 5B). This is consistent with role of the Dot1L inhibitor in decreasing the transcriptional activity on V region in Rep161 cell line which clearly affects a broad range of transcription associated proteins (Fig. 5B). Other proteins may be recruited to the V region perhaps to compensate for the function of the proteins/pathways that were decreased on the V region after inhibitor treatment. Amongst these, 13 proteins are involved in the “RNA helicase activity” pathway (Fig. 5C that are involved in the processing and shaping of RNA molecules especially for ribosome biosynthesis including spliceosome assembly and transcriptional elongation and binding to G4 structures. One of the most enriched proteins in the inhibitor treated V region is DHX36 (Fig. 5C) which is a dominant resolvase in human cells and binds G4-nucleic acids with a high affinity and unwinds them (Yangyuoru et al., 2018). G4-structures also bind AID and are thought to play a role in SHM (Tang and MacCarthy, 2021; Yu et al., 2021a).

**Figure 5.**
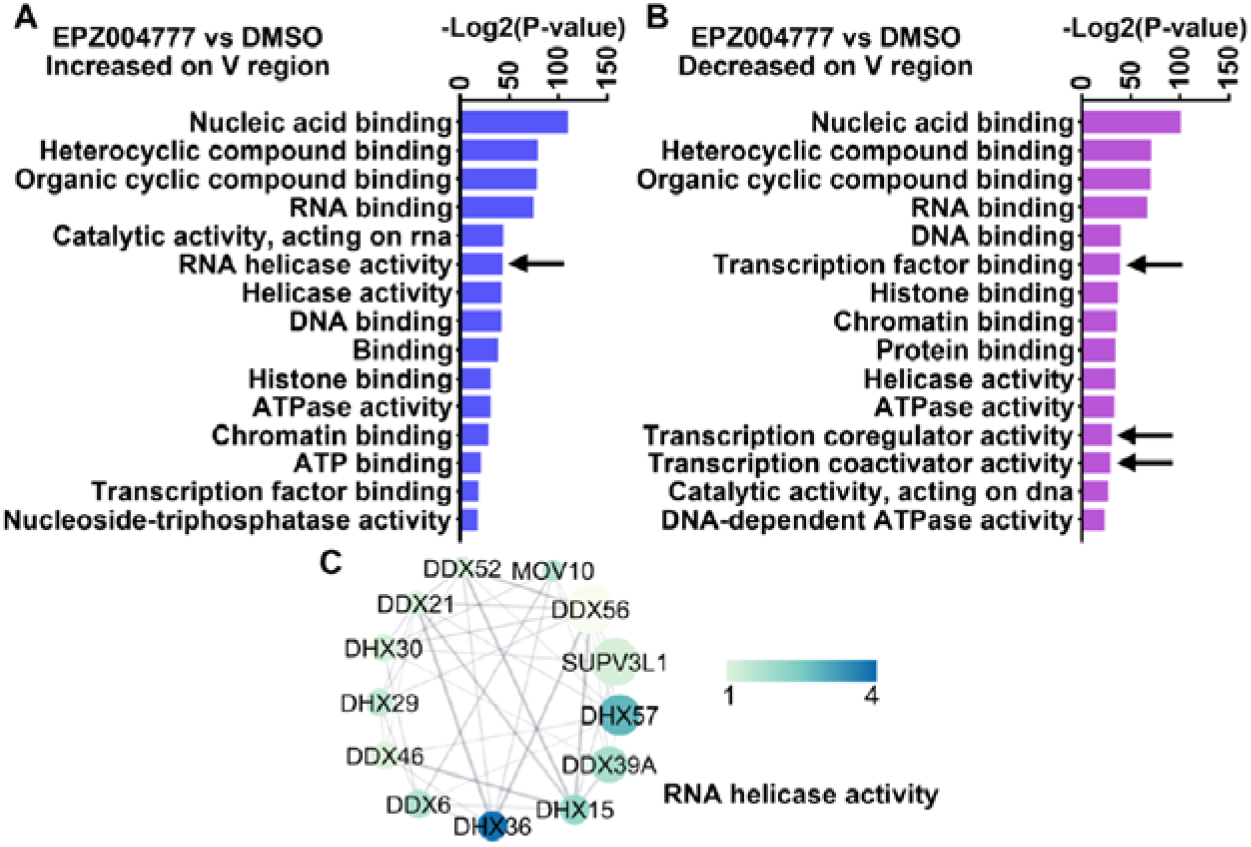
Functional annotation of proteins whose abundance changes on the V region after Dot1L inhibitor treatment. (A-B) Top 15 classifications of molecular function for proteins that are enriched and decreased on V region after EPZ004777 treatment, respectively. The annotation groups that are indicated by black arrow are significantly enriched in each group. (C) Functional network analysis by STRING of proteins that are associated with RNA helicase activity and significantly enriched on the V region. The size of the circle represents the P-value with bigger circles representing more significance. The color indicated by the color band represents the fold change of each protein by comparing its abundance in EPZ treated group with the DMSO control.

### Identification of proteins that are significantly changed on constant Cμ region after EPZ004777 treatment and their functional annotation

We also used the combination of dCas9-APEX and EPZ004777 inhibitor treatment to examine the C-gRNA cell line to study the dynamic protein context of the C region chromatin. By ChIP-qPCR, EPZ004777 treatment significantly reduced the abundance of both H3K79me2/3 and H3K79me1 on the C region while it did not affect the abundance of total H3 (Fig. S2F-S2H). The increase in H3K79me1 in the V region but not the C region with inhibitor treatment was reported in our previous paper (Duan et al., 2021). We found that the EPZ004777 treatment significantly but very slightly decreased the abundance of dCas9-APEX on chromatin in C-gRNA cells after the inhibitor treatment (Fig. S3B). Since the relative abundance of H3K79me 2/3 and H3K79me1 in inhibitor treated cells was similar to what we had observed in non-dCas9-APEX expressing cells, we are assuming that this difference did not significantly affect the relative efficiency of the biotinylation. By differential comparative analysis, EPZ004777 treatment significantly upregulated the abundance of 662 proteins while downregulating the occupancy of 437 proteins on the C region (Fig. 6A). Among the upregulated proteins, we found the transcription elongation factor CDK9 and RNA splicing factor CWC15 (Fig. 6B-C). Among the downregulated proteins, we identified MacroH2A1 (H2AFY), which as noted above is a variant H2A involved in transcription and DNA repair, and NSD2, which dimethylates the histone H3 residue H3K36 (Fig. 6D-E).

**Figure 6.**
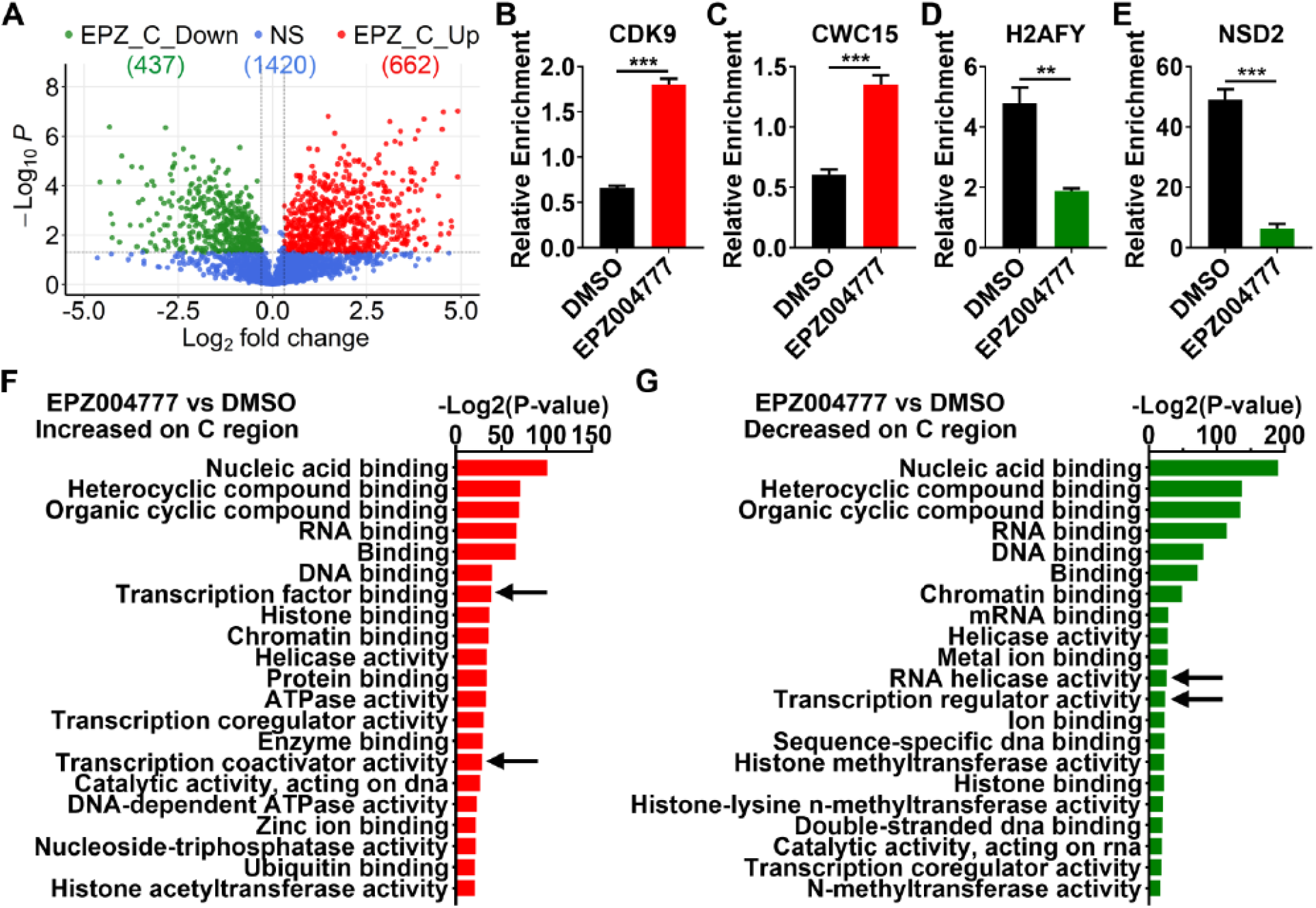
Quantitative and functional analysis of protein change on the C region after Dot1L inhibitor treatment. (A) Volcano plot showing the fold change comparison of proteins on C region between EPZ treatment and DMSO control with p-value distribution. (B-E) Representative proteins with significant changes of their respective relative abundance by Mass Spectrometry on the C region after EPZ treatment. Error bars represent Standard Error of the Mean (SEM) of three independent Mass Spectrometry determinations. (F-G) Top 21 classifications of molecular function for proteins that are enriched and decreased on C region after EPZ004777 treatment, respectively. The annotation groups that are indicated by black arrow are exclusively enriched in each group.

There were many other differentially changed proteins and these were subjected to “Molecular Function” GO annotation (Fig. 6 F and G). Members of these functional groups are both increased and decreased in the C region after inhibitor treatment although some processes are different as indicated by the black arrows. The important point is that many of the dynamic changes in the C region are different from the changes on the V region after EPZ004777. For example, the “RNA helicase activity” is increased on the V region and “transcription related” and “mRNA splicing” factors are decreased on V region (Fig. 5), suggesting that the transcription and RNA metabolism may be regulated differently and through different mechanisms in V and C and that the EPZ004777 inhibitor treatment also affects the V and C regions differently. In order to even begin to understand how these changes affect transcriptional pausing, backtracking and elongation that are thought to facilitate AID mutation, each of these findings will have to be dissected in detail which could be done using inhibitors of other key factors. For example, such studies might explain the different effects of the inhibitor on H3K79me1 at the V and C regions seen in Fig. S2. Since the analysis of impact of the inhibitor on the chromatin context by Mass Spectrometry reveals so many factors that have not been considered before, it offers an opportunity to gain new insights but it also reveals the complexity of these dynamic processes.

## Discussion

The AID-dependent hypermutation of variable and switch regions in antibody genes is a tightly regulated and complicated process. Although proteomic investigations of AID interacting proteins have revealed many factors involved in AID-induced mutation, except for the deletion of AID, the loss of any one of these factors does not eliminate SHM of the V region, confirming that there must be multiple overlapping pathways that are involved in SHM (Methot and Di Noia, 2017). Furthermore, AID is the only factor involved in SHM that is greatly enriched in B cells and all of the other factors identified so far carry out functions genome wide. Most importantly, none of the other factors have so far explained how AID-induced mutations are so highly targeted to the IgV and S regions. Recent studies have shown that the V region is part of a topologically associated chromatin domain (TAD) and a collection of cis-acting DNA sequences called DIVACS contribute to the targeting of AID induce deamination (Senigl et al., 2019), although it has been difficult to identify particular DNA sequences and factors that are responsible for the very high frequency of AID induced mutations. While it would be useful to do a genome-wide screen for V region mutation, this is likely to reveal many indirect effects. An alternative and more direct approach is to examine the relative abundance of proteins that are present in the chromatin of the V region and compare them to other highly transcribed but non-mutating sites such as the C region which is nearby in the Ig locus. This would be even more useful if changes in the relative abundance of these proteins was also examined when specific factors or pathways were interfered with for example with inhibitors. The use of such inhibitors has the advantage of relatively short term treatments and also eliminates the problem of clonal variation if this was being studied in cells in culture. It has now become possible to do this using dCas9-APEX and Mass Spectrometry, or other related techniques (Gauchier et al., 2020), that biotin label the macromolecules in situ in live cells at specific short DNA regions with high resolution. It is especially important that this approach provides a relatively quantitative readout for hundreds of proteins and that the chromatin-associated factors are labeled in a relatively unbiased manner so that new factors and pathways can be discovered.

In the current study, we implemented the dCas9-APEX technique in the Rep161 human Ramos B cell line to compare the chromatin proteomes in V and the C region. The results demonstrate that this dCas9-APEX technique can be used to distinguish genomic loci that are only separated by 8KB in a single gene. Due to the lack of antibodies that can be used for ChIP for most proteins that are differentially distributed and identified by Mass Spectrometry, we could only perform ChIP-qPCR to confirm a few of these factors. As noted in the results, while the trends of the relative abundancies of some of these proteins by ChIP-qPCR in V vs C were similar to the Mass Spectrometry result, the quantifications were often quite different (Figs. 2, 3, 4 and S2). This was expected since ChIPs were done on cross-linked chromatin that had been subjected to a different treatment and also depends on the affinities of the antibodies and the accessibility of the epitopes and no doubt other factors. These results are encouraging and further prove that the dCas9-APEX technique allows us to identify many of the proteins on an interesting genomic locus.

The validity of the dCas9-APEX technique was further confirmed by the identification of many of the proteins that have been individually shown to play roles in SHM, like CTNNBL1, SSRP1, HIRA and Spt5. Moreover, the potential power of this technique was illustrated by the finding that some of these factors, while enriched in the heavy chain locus, were present in approximately the same abundance on both the mutating V region and the non-mutating C region. This suggests that each of these particular proteins is not sufficient by themselves to make ssDNA in the V region accessible to AID induce deamination. In fact, AID itself had a similar abundance by Mass Spectrometry on the V and C regions while Spt5, which directly interacts with AID, was enriched in the V region and not the C region (Table S1). Spt5 has been shown to associate with the V region even in the absence of AID (Methot et al., 2018) so perhaps this is not surprising. However, the many changes that were observed locally when Dot1L activity was only partially inhibited reveal how difficult it will be to demonstrate direct effects. Another interesting finding is that BRAC1 related repair proteins (BRAP, BRAT1 and BARD1) are enriched on the C region raising the possibility that the C region might be mutated by AID but repaired with high fidelity. It is believed that as many as 50% of the AID mutations in the V region are also repaired with high fidelity (Feng et al., 2020), so it is certainly worth investigating more rigorously if the C region is mutated especially in cells lacking some of the repair factors we have found there. Some transcription elongation and RNA splicing factors that have been shown to contribute to V region hypermutation are also found to be enriched on the V region, raising the possibility that has been discussed by others that they are involved in the final “licensing” of AID mutation (Methot et al., 2018; Sun et al., 2013). Since the V region exon does not have any introns except for the one between the leader and framework 1 coding sequences and at the 3’ end of the exon, it is not clear exactly how these splicing factors contribute to SHM but an examination of that is now be more likely to be successful since we have identified some previously unrecognized factors such as ISY1 that are significantly enriched in the V region (Fig.2). While additional detailed studies will be required to better understand the exact role of each of these factors and the complexes they form, the difficulty in establishing roles for at least some of these factors is illustrated by CWC15 which seemed a good candidate to study because of its very low level on the C region and relatively high abundance on both the V and switch μ (Sμ) regions (Figure 2A and D) which both undergo AID induced mutations. Unluckily, the KO of CWC15 is lethal to the Ramos cells suggesting not surprisingly that CWC15 may be very important not just on the V region but genome-wide and its further study will require very short-term conditional interference in Ramos and other B cells.

The Dot1L inhibitor revealed that changes of protein pathways on V region after Dot1L inhibitor treatment are different from the C region (Fig. 5 and 6), suggesting the protein complexes associated with V region that regulate its transcription and SHM are different from the C region that does not have detectable SHM. Encouragingly, the abundance of MLLT1, a component of super elongation complex, was increased on V region after Dot1L inhibitor treatment. We had shown previously that the Dot1L inhibitor EPZ004777 only inhibited its enzymatic activity but did not change its physical abundance on the V region, and the EPZ004777 treatment only decreased the SHM by 30% compared with the 70% decrease of SHM in Dot1L KO cells (Duan et al., 2021). The Dot1L molecule has been shown to both form its own complex and to be a component of SEC where its function appears to be independent of its histone methyltransferase activity (Cao et al., 2020). We found that the SEC inhibitor KL-2 can decrease the V region mutation in our Rep161 and the combination of even low doses of the Dot1L inhibitor and the KL-2 significantly decreased SHM to a greater degree than the sum of their single treatments (p = 0.03). This suggests that even though Dot1L can participate in the SEC complex, it is playing an independent role at the V region and confirms our impression that multiple elongation complexes are required for AID-induced mutation and play different roles. In the current study we have not used dCas9-APEX to study the difference of combinatorial histone modifications between V and C regions. This is an interesting direction that should be pursued in the future. Also, with this technique, the protein complexes on the 3’ regulatory region (3’ RR) of the IgH locus that plays important roles in SHM and CSR by looping can also be explored. Obviously, this technique can also be transferred to other biological questions such as identifying proteins involved into gene translocation and identifying how certain genes are actively transcribed even though they lack the typical markers (e.g. H3K4me3, H3K27ac).

## Materials and Methods

### Cell line culture

The Reporter 161 (Rep161) cell line was generated using the Ramos human Burkitt’s lymphoma B cell line as reported previously (Duan et al., 2021; Wang et al., 2014). Rep161 was cultured with Iscove’s modified Dulbecco’s medium (IMDM, BioWhittaker) supplemented with 10% of the fetal bovine serum and Penicillin-Streptomycin. In order to stimulate SHM, Rep161 cells were treated with 0.25 μM 4-hydroxytamoxifen (4-OHT) for 7 days and then were analyzed by flow cytometry to determine the frequency of cells with a decrease in mCherry fluorescence, which represents the frequency of mutation by AID.

### dCas9-APEX transfection

The doxycycline inducible dCas9-APEX vector was purchased from Addgene (#97421) and electroporated into Rep161 cells. 2 days after electroporation, 0.4 μg/ml puromycin was added to the culture to obtain pools of stable puromycin resistant cells that were plated out in limiting dilution in 30 96-well plates. Clones that expressed the dCas9-APEX protein after doxycycline treatment were identified visually and merged together as a pool that was used for all of the following experiments. The pooled clones were maintained in 0.2 μg/ml puromycin to maintain expression of dCas9-APEX.

### Guide RNA (gRNA) design and infection

The Benchling (https://www.benchling.com/crispr/) website was used to design gRNA targeting the V and C regions, separately. Based on the criteria of high efficiency of targeting and low chance of off-targeting, the sequence of the gRNA targeting V region is 5’-GGAACAATACGAACGAGCCG-3’ and for C region is 5’-CCACTGCACGAAGACGTCCG-3’. To achieve a relatively higher efficiency of transfection for these gRNAs, a 3^rd^ generation of lenti virus expression vector with the selection marker for blasticidin S deaminase (BSD) was prepared for each gRNA. Lenti particles were prepared for each gRNA and added to two populations of stable dCas9-APEX expressing cells, respectively. After two days, 10 μg/ml of Blasticidin S HCl was added to each transfection to obtain the stable cell lines. After obtaining the stable positive clones, 0.2 μg/ml puromycin and 5 μg/ml of Blasticidin S HCl were used in the culture to maintain expression of dCas9-APEX and the gRNA, separately. The final two stable cell lines were named “V-gRNA” which expressed the dCas9-APEX and gRNA targeted to V region and “C-gRNA” that expressed dCas9-APEX and gRNA targeted to C region.

### Western Blotting

To detect Flag-tagged dCas9-APEX in the whole cell lysate, cells were collected and rinsed with ice cold PBS twice before adding RIPA buffer containing a protein inhibitor cocktail. The cells were then incubated at 4 °C with gentle rotation for 30min and centrifuged at 13,000 RPM for 10 min to collect the supernatant for WB using Anti-Flag antibody (F1804, Sigma). For the WB using chromatin after the biotinylation reaction, after RIPA buffer incubation the tube was centrifuged with 10,000 PRM for 5min, the pellet was gently suspended with 1 ml of RIPA buffer, and the tube was centrifuged again to obtain the chromatin pellet. The chromatin was resuspended with RIPA buffer containing 0.5% SDS and sonicated using Bioruptor for 2 cycles (30 seconds ON/30 seconds OFF, 5min for each cycle with power set to High) at 4°C and then centrifuged and the soluble chromatin fraction was used for WB (Yu et al., 2021b). The information about the antibodies used for WB is listed in Supplementary Table S3.

### Biotinylation reaction and pull down

Due to the very low transfection and expression of ecotopic genes in Ramos, to get adequate expression levels of dCas9-APEX, the V-gRNA and C-gRNA cell lines were treated with 0.2ug/ml Doxycycline for 3 days, and then 40 million cells for each cell line was collected to prepare 8ml of cells in medium with 5 million cells/ml. Biotin-Tyramide Phenol (dissolved in DMSO, CataLog: LS-3500, Iris Biotech) was added to each cell sample at a final concentration of 500μM and incubated at 37 °C with gentle rotation for 30min. Then a freshly prepared H2O2 solution was added to a final concentration of 1mM and incubated at room temperature for exactly 60 seconds before adding 30ml of ice cold quencher solution (5 mM trolox, 10 mM sodium ascorbate, and 10 mM sodium azide) to stop the reaction. The cells were washed five times (three quencher washes and two PBS washes) to continue the quench and to remove excess BP. Then RIPA buffer containing 0.5% Triton X-100 was added to each cell sample to lyse the cells and the chromatin fraction was collected as described above in the “Western Blotting” section. 1 mg of chromatin fraction was incubated with 80 ul of Streptavidin Dynabeads (CataLog: 65602, Invitrogen) for 3 hours at 4°C with gentle rotation. Then each sample was washed with a series of buffers to remove nonspecifically bound proteins: twice with RIPA lysis buffer, once with 1 M KCl, once with 0.1 M Na_2_CO_3_, once with 2 M urea in 10 mM Tris-HCl, pH 8.0, and twice with RIPA lysis buffer. Proteins were eluted in 30 μl of 3 X protein loading buffer supplemented with 2 mM biotin and 20 mM DTT with heating for 10 min at 100°C and then the pull-down fraction was transferred to a new tube using a magnetic rack. Three immunoprecipitations were performed each time for each cell line in order to have enough pull-down samples and pooled. For each cell line, three independent experiments were performed using the whole process.

### Protein analysis via liquid chromatography coupled to Mass Spectrometry

Proteins were eluted from the immunoprecipitation using a buffer containing 5% SDS, 5 mM DTT and 50 mM ammonium bicarbonate (pH = 8), and left on the bench for about 1 hour for disulfide bond reduction. Samples were then alkylated with 20 mM iodoacetamide in the dark for 30 minutes. Afterward, phosphoric acid was added to the sample at a final concentration of 1.2%. Samples were diluted in six volumes of binding buffer (90% methanol and 10 mM ammonium bicarbonate, pH 8.0). After gentle mixing, the protein solution was loaded to an S-trap filter (Protifi) and spun at 500 g for 30 sec. The sample was washed twice with binding buffer. Finally, 1 μg of sequencing grade trypsin (Promega), diluted in 50 mM ammonium bicarbonate, was added into the S-trap filter and samples were digested at 37 °C for 18 h. Peptides were eluted in three steps: (i) 40 μl of 50 mM ammonium bicarbonate, (ii) 40 μl of 0.1% trifluoroacetic acid (TFA) and (iii) 40 μl of 60% acetonitrile and 0.1% TFA. The peptide solution was pooled, spun at 1,000 g for 30 sec and dried in a vacuum centrifuge (HaileMariam et al., 2018).

Before Mass Spectrometry analysis, samples were desalted using a 96-well plate filter (Orochem) packed with 1 mg of Oasis HLB C-18 resin (Waters). Briefly, the samples were resuspended in 100 μl of 0.1% TFA and loaded onto the HLB resin, which was previously equilibrated using 100 μl of the same buffer. After washing with 100 μl of 0.1% TFA, the samples were eluted with a buffer containing 70 μl of 60% acetonitrile and 0.1% TFA and then dried in a vacuum centrifuge.

Samples were then resuspended in 10 μl of 0.1% TFA and loaded onto a Dionex RSLC Ultimate 3000 (Thermo Scientific) nano-liquid chromatographer coupled online with an Orbitrap Fusion Lumos mass spectrometer (Thermo Scientific). Chromatographic separation was performed with a two-column system, consisting of a C-18 trap cartridge (300 μm ID, 5 mm length) and a picofrit analytical column (75 μm ID, 25 cm length) packed in-house with reversed-phase Repro-Sil Pur C18-AQ 3 μm resin. To analyze the proteome, peptides were separated using a 45 min gradient from 4-30% buffer B (buffer A: 0.1% formic acid, buffer B: 80% acetonitrile + 0.1% formic acid) at a flow rate of 300 nl/min. The mass spectrometer was set to acquire spectra in a data-dependent acquisition (DDA) mode. Briefly, the full MS scan was set to 300-1200 m/z in the orbitrap with a resolution of 120,000 (at 200 m/z) and an AGC target of 5×10e5. MS/MS was performed in the ion trap using the top speed mode (2 secs), an AGC target of 1×10e4 and an HCD collision energy of 35 (Meyer, 2021).

Proteome raw files were searched using Proteome Discoverer software (v2.4, Thermo Scientific) using SEQUEST search engine and the SwissProt human database (updated February 2020) (Diament and Noble, 2011). The search for total proteome included variable modification of N-terminal acetylation, and fixed modification of carbamidomethyl cysteine. Trypsin was specified as the digestive enzyme with up to 2 missed cleavages allowed. Mass tolerance was set to 10 pm for precursor ions and 0.2 Da for product ions. Peptide and protein false discovery rate was set to 1%. Following the search, data was processed as previously described (Aguilan et al., 2020). Briefly, proteins were log2 transformed, normalized by the average value of each sample and missing values were imputed using a normal distribution 2 standard deviations lower than the mean. Statistical regulation was assessed using heteroscedastic T-test (if p-value < 0.05). Data distribution was assumed to be normal but this was not formally tested.

### Functional analysis of the proteomic data

Cytoscape was used for the functional annotation of the MASS Spectrometry data. For functional interaction analysis, STRING in Cytoscape (version 3.8.2) was adopted with cutoff being 0.4. For the Gene Ontology (GO) enrichment analysis, the STRING Enrichment in Cytoscape was used.

### Chromatin immunoprecipitation (ChIP) and qPCR

ChIP was performed as described previously (Yu et al., 2021b). Briefly, 10 million healthy cells were collected in medium and fixed with 1% formaldehyde for 10 min at room temperature and 0.125 M glycine was used to quench the reaction. Then the cells were suspended in 1% SDS lysing buffer and sonicated using Bioruptor to yield DNA fragments between 300bp and 1 kb. 200 μg of chromatin was used for each ChIP. The antibodies and primers used for ChIP were listed in Supplementary Table S2 and Table S3.

### Inhibitor treatment

For the Dot1L inhibitor EPZ004777 treatment, the V-gRNA and C-gRNA cell lines were treated with EPZ004777 (20 μM) plus 0.25 μM 4-OHT for 7 days. Then the cells were collected for SHM analysis by FACS. For the combinational treatment, EPZ004777 (5 μM) and the inhibitor for super elongation complex KL-2 (0.5 μM) were used for 7 days before the SHM analysis by FACS.

### Statistical analysis

The unpaired Student’s T-test from Graphpad Prism 8 was used for all of the studies. P values < 0.05 were considered statistically significant and all error bars represent the standard error of the mean from three independent experiments. For the p-value calculation of the functional enrichment of the MASS Spectrometry dataset, the internal algorithm from STRING was automatically adopted.

## Data availability

Mass spectrometry data of the dCAS proteomics pull-down are deposited on the repository Chorus (https://chorusproject.org) under project number 1756.

## Supplementary material

**Figure S1. The verification of proteins that are not significantly changed between V and C regions by ChIP-qPCR. (A)** Volcano plot showing the fold change comparison of proteins on V and C region with p-value distribution. Representative unchanged proteins were labeled with their respective names. **(B)** ChIP-qPCR verification of proteins that are not changed between V and C regions. RBBP4 is a chromatin remodeling factor; RUVBL2 is DNA helicase; MCM5 and MCM3 are involved in DNA replication. Error bars in this experiment and in the subsequent experiments represent Standard Error of the Mean (SEM) of three independent Mass Spectrometry determinations. * means p-value <= 0.05, ** means p-value <= 0.01, *** means p-value <= 0.001, ns means no significance.

**Figure S2. Dot1L inhibitor treatment inhibited the SHM by decreasing the abundance of H3K79me2/3 on the Ig locus.** (A-B) The effect of Dot1L inhibitor EPZ004777 (20) on SHM of V region in V-gRNA cell line measured by flow cytometry (FACS). The V-gRNA cell line was treated with EPZ004777 (20 μM) plus 4-OHT (0.25 μM) for 7 days and then the percentage of cells with loss of mCherry was measured by FACS to determine the frequency the SHM on V region. (C-E) The effect of EPZ004777 treatment on the abundance of H3K79me2/3, H2K79me1 and H3 on V region in V-gRNA cell line. Cells were treated with 7 days of EPZ004777 (20 μM) plus 4-OHT (0.25 μM) and then were collected for ChIP-qPCR. (F-H) The effect of EPZ004777 treatment on the abundance of H3K79me2/3, H2K79me1 and H3 on C region in C-gRNA cell line. C-gRNA cells were treated with 7 days of EPZ004777 (20 μM) plus 4-OHT (0.25 μM) and then were collected for ChIP-qPCR.

**Figure S3. The effect of various inhibitor treatments on the abundance of dCas9-APEX on chromatin in both V-gRNA and C-gRNA cell lines and the SHM, separately.** (A-B), Y-axis means the normalized dCas9-APEX protein abundance in Log2 fold from the Mass Spectrometry. (C), representative FACS data of 3 independent repeats related to Fig. 4L showed the effect of Dot1L inhibitor EPZ004777 (5 μM), super elongation complex inhibitor KL-2 (0.5 μM), and the combination of EPZ004777 (5 μM) and KL-2 (0.5 μM) on SHM.

**Table S1. The relative abundance comparison of proteins identified between V and C regions that are known to play roles in SHM in B cells.**

**Table S2. The information on the primers used for ChIP-qPCR in this study.**

**Table S3. The information on the antibodies used in this study.**

## Author Contributions

G. Yu performed most of the experiments, analyzed the data and wrote the first draft of the manuscript. Z. Duan performed the inhibitor treatment experiments and Y. Zhang constructed the vectors. J. Aguilan participated in the Mass Spectrometry. G. Yu, M. D. Scharff and S. Sidoli designed the study, interpreted data and revised the manuscript.

## Acknowledgments

We thank the core facilities at Albert Einstein College of Medicine for their technical support for flow cytometry and Mass Spectrometry. Funding for this work was provided by NIH Grant R01 AI132507-01A1 to MDS. G.Y. was supported in part by the American Association of Immunologists Intersect Fellowship. The Sidoli lab gratefully acknowledges the Einstein-Mount Sinai Diabetes Research Center (Pilot Grant), AFAR (Sagol Network GerOmics award), Deerfield (Xseed award), Relay Therapeutics, Merck and the NIH Office of the Director (1S10OD030286-01). The funders had no role in study design, data collection and analysis, decision to publish, or preparation of the manuscript.

## References

Aguilan, J.T., K. Kulej, and S. Sidoli. 2020. Guide for protein fold change and p-value calculation for non-experts in proteomics. Mol Omics 16:573–582.

Aida, M., N. Hamad, A. Stanlie, N.A. Begum, and T. Honjo. 2013. Accumulation of the FACT complex, as well as histone H3.3, serves as a target marker for somatic hypermutation. Proc Natl Acad Sci U S A 110:7784–7789.

Álvarez-Prado Á, F., P. Pérez-Durán, A. Pérez-García, A. Benguria, C. Torroja, V.G. de Yébenes, and A.R. Ramiro. 2018. A broad atlas of somatic hypermutation allows prediction of activation-induced deaminase targets. J Exp Med 215:761–771.

Ashburner, M., C.A. Ball, J.A. Blake, D. Botstein, H. Butler, J.M. Cherry, A.P. Davis, K. Dolinski, S.S. Dwight, J.T. Eppig, M.A. Harris, D.P. Hill, L. Issel-Tarver, A. Kasarskis, S. Lewis, J.C. Matese, J.E. Richardson, M. Ringwald, G.M. Rubin, and G. Sherlock. 2000. Gene ontology: tool for the unification of biology. The Gene Ontology Consortium. Nat Genet 25:25–29.

Basu, U., F.L. Meng, C. Keim, V. Grinstein, E. Pefanis, J. Eccleston, T. Zhang, D. Myers, C.R. Wasserman, D.R. Wesemann, K. Januszyk, R.I. Gregory, H. Deng, C.D. Lima, and F.W. Alt. 2011. The RNA exosome targets the AID cytidine deaminase to both strands of transcribed duplex DNA substrates. Cell 144:353–363.

Bransteitter, R., P. Pham, M.D. Scharff, and M.F. Goodman. 2003. Activation-induced cytidine deaminase deaminates deoxycytidine on single-stranded DNA but requires the action of RNase. Proc Natl Acad Sci U S A 100:4102–4107.

Cao, K., M. Ugarenko, P.A. Ozark, J. Wang, S.A. Marshall, E.J. Rendleman, K. Liang, L. Wang, L. Zou, E.R. Smith, F. Yue, and A. Shilatifard. 2020. DOT1L-controlled cell-fate determination and transcription elongation are independent of H3K79 methylation. Proc Natl Acad Sci U S A 117:27365–27373.

Casellas, R., U. Basu, W.T. Yewdell, J. Chaudhuri, D.F. Robbiani, and J.M. Di Noia. 2016. Mutations, kataegis and translocations in B cells: understanding AID promiscuous activity. Nat Rev Immunol 16:164–176.

Chen, C.C., W. Feng, P.X. Lim, E.M. Kass, and M. Jasin. 2018a. Homology-Directed Repair and the Role of BRCA1, BRCA2, and Related Proteins in Genome Integrity and Cancer. Annu Rev Cancer Biol 2:313–336.

Chen, F.X., E.R. Smith, and A. Shilatifard. 2018b. Born to run: control of transcription elongation by RNA polymerase II. Nat Rev Mol Cell Biol 19:464–478.

Chen, J., Z. Cai, M. Bai, X. Yu, C. Zhang, C. Cao, X. Hu, L. Wang, R. Su, D. Wang, L. Wang, Y. Yao, R. Ye, B. Hou, Y. Yu, S. Yu, J. Li, and Y. Xue. 2018c. The RNA-binding protein ROD1/PTBP3 cotranscriptionally defines AID-loading sites to mediate antibody class switch in mammalian genomes. Cell Res 28:981–995.

Deshpande, A.J., A. Deshpande, A.U. Sinha, L. Chen, J. Chang, A. Cihan, M. Fazio, C.W. Chen, N. Zhu, R. Koche, L. Dzhekieva, G. Ibáñez, S. Dias, D. Banka, A. Krivtsov, M. Luo, R.G. Roeder, J.E. Bradner, K.M. Bernt, and S.A. Armstrong. 2014. AF10 regulates progressive H3K79 methylation and HOX gene expression in diverse AML subtypes. Cancer Cell 26:896–908.

Diament, B.J., and W.S. Noble. 2011. Faster SEQUEST searching for peptide identification from tandem mass spectra. J Proteome Res 10:3871–3879.

Dinesh, R.K., B. Barnhill, A. Ilanges, L. Wu, D.A. Michelson, F. Senigl, J. Alinikula, J. Shabanowitz, D.F. Hunt, and D.G. Schatz. 2020. Transcription factor binding at Ig enhancers is linked to somatic hypermutation targeting. Eur J Immunol 50:380–395.

Duan, Z., L.B. Baughn, X. Wang, Y. Zhang, V. Gupta, T. MacCarthy, M.D. Scharff, and G. Yu. 2021. Role of Dot1L and H3K79 methylation in regulating somatic hypermutation of immunoglobulin genes. Proceedings of the National Academy of Sciences 118:e2104013118.

Feng, Y., C. Li, J.A. Stewart, P. Barbulescu, N. Seija Desivo, A. Álvarez-Quilón, R.C. Pezo, M.L.W. Perera, K. Chan, A.H.Y. Tong, R. Mohamad-Ramshan, M. Berru, D. Nakib, G. Li, G.A. Kardar, J.R. Carlyle, J. Moffat, D. Durocher, J.M. Di Noia, A.S. Bhagwat, and A. Martin. 2021. FAM72A antagonizes UNG2 to promote mutagenic repair during antibody maturation. Nature 600:324–328.

Feng, Y., N. Seija, J.M. Di Noia, and A. Martin. 2020. AID in Antibody Diversification: There and Back Again. Trends Immunol 41:586–600.

Friedl, E.M., W.S. Lane, H. Erdjument-Bromage, P. Tempst, and D. Reinberg. 2003. The C-terminal domain phosphatase and transcription elongation activities of FCP1 are regulated by phosphorylation. Proc Natl Acad Sci U S A 100:2328–2333.

Ganesh, K., S. Adam, B. Taylor, P. Simpson, C. Rada, and M. Neuberger. 2011. CTNNBL1 is a novel nuclear localization sequence-binding protein that recognizes RNA-splicing factors CDC5L and Prp31. J Biol Chem 286:17091–17102.

Gao, X.D., L.C. Tu, A. Mir, T. Rodriguez, Y. Ding, J. Leszyk, J. Dekker, S.A. Shaffer, L.J. Zhu, S.A. Wolfe, and E.J. Sontheimer. 2018. C-BERST: defining subnuclear proteomic landscapes at genomic elements with dCas9-APEX2. Nat Methods 15:433–436.

Garrett, K.P., T. Aso, J.N. Bradsher, S.I. Foundling, W.S. Lane, R.C. Conaway, and J.W. Conaway. 1995. Positive regulation of general transcription factor SIII by a tailed ubiquitin homolog. Proc Natl Acad Sci U S A 92:7172–7176.

Gauchier, M., G. van Mierlo, M. Vermeulen, and J. Déjardin. 2020. Purification and enrichment of specific chromatin loci. Nat Methods 17:380–389.

HaileMariam, M., R.V. Eguez, H. Singh, S. Bekele, G. Ameni, R. Pieper, and Y. Yu. 2018. S-Trap, an Ultrafast Sample-Preparation Approach for Shotgun Proteomics. J Proteome Res 17:2917–2924.

Jeevan-Raj, B.P., I. Robert, V. Heyer, A. Page, J.H. Wang, F. Cammas, F.W. Alt, R. Losson, and B. Reina-San-Martin. 2011. Epigenetic tethering of AID to the donor switch region during immunoglobulin class switch recombination. J Exp Med 208:1649–1660.

Kobayashi, M., M. Aida, H. Nagaoka, N.A. Begum, Y. Kitawaki, M. Nakata, A. Stanlie, T. Doi, L. Kato, I.M. Okazaki, R. Shinkura, M. Muramatsu, K. Kinoshita, and T. Honjo. 2009. AID-induced decrease in topoisomerase 1 induces DNA structural alteration and DNA cleavage for class switch recombination. Proc Natl Acad Sci U S A 106:22375–22380.

Liang, K., E.R. Smith, Y. Aoi, K.L. Stoltz, H. Katagi, A.R. Woodfin, E.J. Rendleman, S.A. Marshall, D.C. Murray, L. Wang, P.A. Ozark, R.K. Mishra, R. Hashizume, G.E. Schiltz, and A. Shilatifard. 2018. Targeting Processive Transcription Elongation via SEC Disruption for MYC-Induced Cancer Therapy. Cell 175:766–779.e717.

Luo, Z., C. Lin, and A. Shilatifard. 2012. The super elongation complex (SEC) family in transcriptional control. Nat Rev Mol Cell Biol 13:543–547.

Manis, J.P., J.C. Morales, Z. Xia, J.L. Kutok, F.W. Alt, and P.B. Carpenter. 2004. 53BP1 links DNA damage-response pathways to immunoglobulin heavy chain class-switch recombination. Nature immunology 5:481–487.

Manning, B.J., and T. Yusufzai. 2017. The ATP-dependent chromatin remodeling enzymes CHD6, CHD7, and CHD8 exhibit distinct nucleosome binding and remodeling activities. J Biol Chem 292:11927–11936.

Methot, S.P., and J.M. Di Noia. 2017. Molecular Mechanisms of Somatic Hypermutation and Class Switch Recombination. Adv Immunol 133:37–87.

Methot, S.P., L.C. Litzler, P.G. Subramani, A.K. Eranki, H. Fifield, A.M. Patenaude, J.C. Gilmore, G.E. Santiago, H. Bagci, J.F. Côté, M. Larijani, R.E. Verdun, and J.M. Di Noia. 2018. A licensing step links AID to transcription elongation for mutagenesis in B cells. Nat Commun 9:1248.

Meyer, J.G. 2021. Qualitative and Quantitative Shotgun Proteomics Data Analysis from Data-Dependent Acquisition Mass Spectrometry. Methods in molecular biology 2259:297–308.

Myers, S.A., J. Wright, R. Peckner, B.T. Kalish, F. Zhang, and S.A. Carr. 2018. Discovery of proteins associated with a predefined genomic locus via dCas9-APEX-mediated proximity labeling. Nat Methods 15:437–439.

Nowak, U., A.J. Matthews, S. Zheng, and J. Chaudhuri. 2011. The splicing regulator PTBP2 interacts with the cytidine deaminase AID and promotes binding of AID to switch-region DNA. Nature immunology 12:160–166.

Oudinet, C., F.Z. Braikia, A. Dauba, J.M. Santos, and A.A. Khamlichi. 2019. Developmental regulation of DNA cytosine methylation at the immunoglobulin heavy chain constant locus. PLoS genetics 15:e1007930.

Parsa, J.Y., S. Ramachandran, A. Zaheen, R.M. Nepal, A. Kapelnikov, A. Belcheva, M. Berru, D. Ronai, and A. Martin. 2012. Negative supercoiling creates single-stranded patches of DNA that are substrates for AID-mediated mutagenesis. PLoS genetics 8:e1002518.

Pavri, R., A. Gazumyan, M. Jankovic, M. Di Virgilio, I. Klein, C. Ansarah-Sobrinho, W. Resch, A. Yamane, B. Reina San-Martin, V. Barreto, T.J. Nieland, D.E. Root, R. Casellas, and M.C. Nussenzweig. 2010. Activation-induced cytidine deaminase targets DNA at sites of RNA polymerase II stalling by interaction with Spt5. Cell 143:122–133.

Peled, J.U., F.L. Kuang, M.D. Iglesias-Ussel, S. Roa, S.L. Kalis, M.F. Goodman, and M.D. Scharff. 2008. The biochemistry of somatic hypermutation. Annual review of immunology 26:481–511.

Pham, P., S. Malik, C. Mak, P.C. Calabrese, R.G. Roeder, and M.F. Goodman. 2019. AID-RNA polymerase II transcription-dependent deamination of IgV DNA. Nucleic Acids Res 47:10815–10829.

Qian, J., Q. Wang, M. Dose, N. Pruett, K.R. Kieffer-Kwon, W. Resch, G. Liang, Z. Tang, E. Mathe, C. Benner, W. Dubois, S. Nelson, L. Vian, T.Y. Oliveira, M. Jankovic, O. Hakim, A. Gazumyan, R. Pavri, P. Awasthi, B. Song, G. Liu, L. Chen, S. Zhu, L. Feigenbaum, L. Staudt, C. Murre, Y. Ruan, D.F. Robbiani, Q. Pan-Hammarstrom, M.C. Nussenzweig, and R. Casellas. 2014. B cell super-enhancers and regulatory clusters recruit AID tumorigenic activity. Cell 159:1524–1537.

Senigl, F., Y. Maman, R.K. Dinesh, J. Alinikula, R.B. Seth, L. Pecnova, A.D. Omer, S.S.P. Rao, D. Weisz, J.M. Buerstedde, E.L. Aiden, R. Casellas, J. Hejnar, and D.G. Schatz. 2019. Topologically Associated Domains Delineate Susceptibility to Somatic Hypermutation. Cell Rep 29:3902–3915.e3908.

Shannon, P., A. Markiel, O. Ozier, N.S. Baliga, J.T. Wang, D. Ramage, N. Amin, B. Schwikowski, and T. Ideker. 2003. Cytoscape: a software environment for integrated models of biomolecular interaction networks. Genome research 13:2498–2504.

Singh, S.K., K. Maeda, M.M. Eid, S.A. Almofty, M. Ono, P. Pham, M.F. Goodman, and N. Sakaguchi. 2013. GANP regulates recruitment of AID to immunoglobulin variable regions by modulating transcription and nucleosome occupancy. Nat Commun 4:1830.

Sun, J., G. Rothschild, E. Pefanis, and U. Basu. 2013. Transcriptional stalling in B-lymphocytes: a mechanism for antibody diversification and maintenance of genomic integrity. Transcription 4:127–135.

Sun, Z., and E. Bernstein. 2019. Histone variant macroH2A: from chromatin deposition to molecular function. Essays Biochem 63:59–74.

Suskiewicz, M.J., F. Zobel, T.E.H. Ogden, P. Fontana, A. Ariza, J.C. Yang, K. Zhu, L. Bracken, W.J. Hawthorne, D. Ahel, D. Neuhaus, and I. Ahel. 2020. HPF1 completes the PARP active site for DNA damage-induced ADP-ribosylation. Nature 579:598–602.

Szklarczyk, D., A.L. Gable, K.C. Nastou, D. Lyon, R. Kirsch, S. Pyysalo, N.T. Doncheva, M. Legeay, T. Fang, P. Bork, L.J. Jensen, and C. von Mering. 2021. The STRING database in 2021: customizable protein-protein networks, and functional characterization of user-uploaded gene/measurement sets. Nucleic Acids Res 49:D605–d612.

Tang, C., and T. MacCarthy. 2021. Characterization of DNA G-Quadruplex Structures in Human Immunoglobulin Heavy Variable (IGHV) Genes. Front Immunol 12:671944.

Tarsalainen, A., Y. Maman, F.L. Meng, M.K. Kyläniemi, A. Soikkeli, P. Budzyńska, J.J. McDonald, F. Šenigl, F.W. Alt, D.G. Schatz, and J. Alinikula. 2022. Ig Enhancers Increase RNA Polymerase II Stalling at Somatic Hypermutation Target Sequences. J Immunol 208:143–154.

van Maldegem, F., S. Maslen, C.M. Johnson, A. Chandra, K. Ganesh, M. Skehel, and C. Rada. 2015. CTNNBL1 facilitates the association of CWC15 with CDC5L and is required to maintain the abundance of the Prp19 spliceosomal complex. Nucleic Acids Res 43:7058–7069.

Victora, G.D., and M.C. Nussenzweig. 2022. Germinal Centers. Annual review of immunology

Voong, C.K., J.A. Goodrich, and J.F. Kugel. 2021. Interactions of HMGB Proteins with the Genome and the Impact on Disease. Biomolecules 11:

Wang, X., M. Fan, S. Kalis, L. Wei, and M.D. Scharff. 2014. A source of the single-stranded DNA substrate for activation-induced deaminase during somatic hypermutation. Nat Commun 5:4137.

Willmann, K.L., S. Milosevic, S. Pauklin, K.M. Schmitz, G. Rangam, M.T. Simon, S. Maslen, M. Skehel, I. Robert, V. Heyer, E. Schiavo, B. Reina-San-Martin, and S.K. Petersen-Mahrt. 2012. A role for the RNA pol II-associated PAF complex in AID-induced immune diversification. J Exp Med 209:2099–2111.

Yangyuoru, P.M., D.A. Bradburn, Z. Liu, T.S. Xiao, and R. Russell. 2018. The G-quadruplex (G4) resolvase DHX36 efficiently and specifically disrupts DNA G4s via a translocation-based helicase mechanism. J Biol Chem 293:1924–1932.

Yu, G., Y. Wu, Z. Duan, C. Tang, H. Xing, M.D. Scharff, and T. MacCarthy. 2021a. A Bayesian model based computational analysis of the relationship between bisulfite accessible single-stranded DNA in chromatin and somatic hypermutation of immunoglobulin genes. PLoS Comput Biol 17:e1009323.

Yu, G., Y. Zhang, V. Gupta, J. Zhang, T. MacCarthy, Z. Duan, and M.D. Scharff. 2021b. The role of HIRA-dependent H3.3 deposition and its modifications in the somatic hypermutation of immunoglobulin variable regions. Proc Natl Acad Sci U S A 118:

