## Supplementary Material for "Locus-specific proteomics identifies new aspects of the chromatin context involved in V region somatic hypermutation"

**Figure S1.**

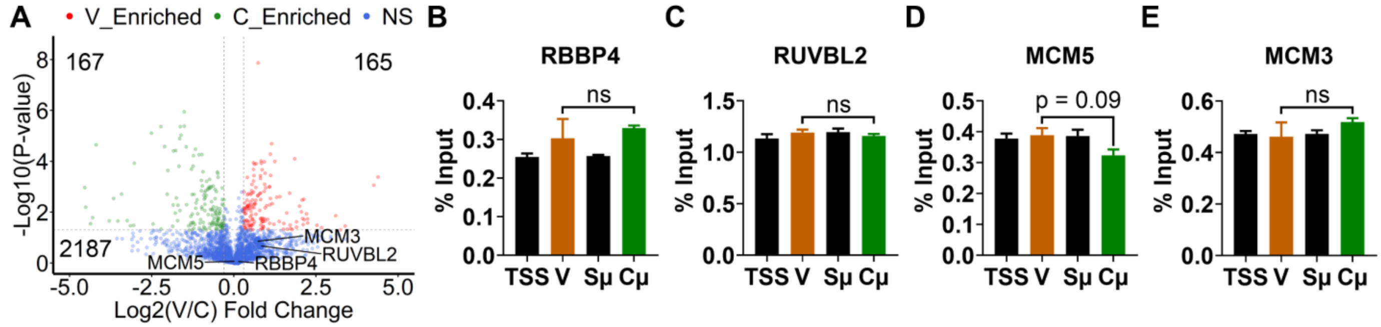


**Figure S2**

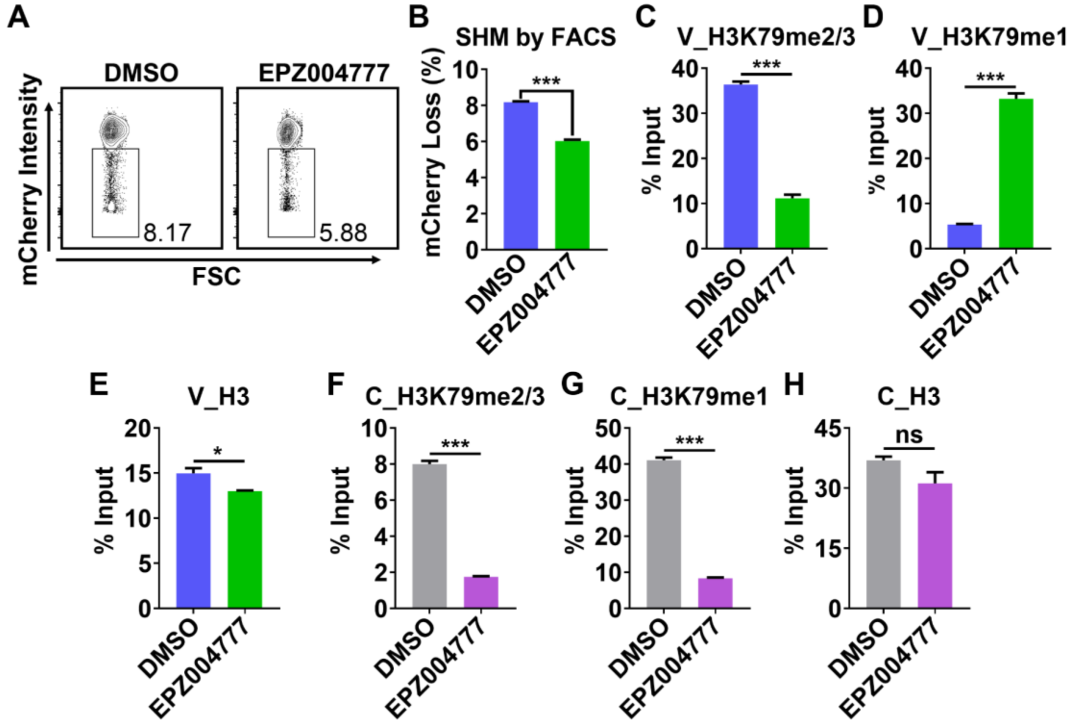


**Figure S3**

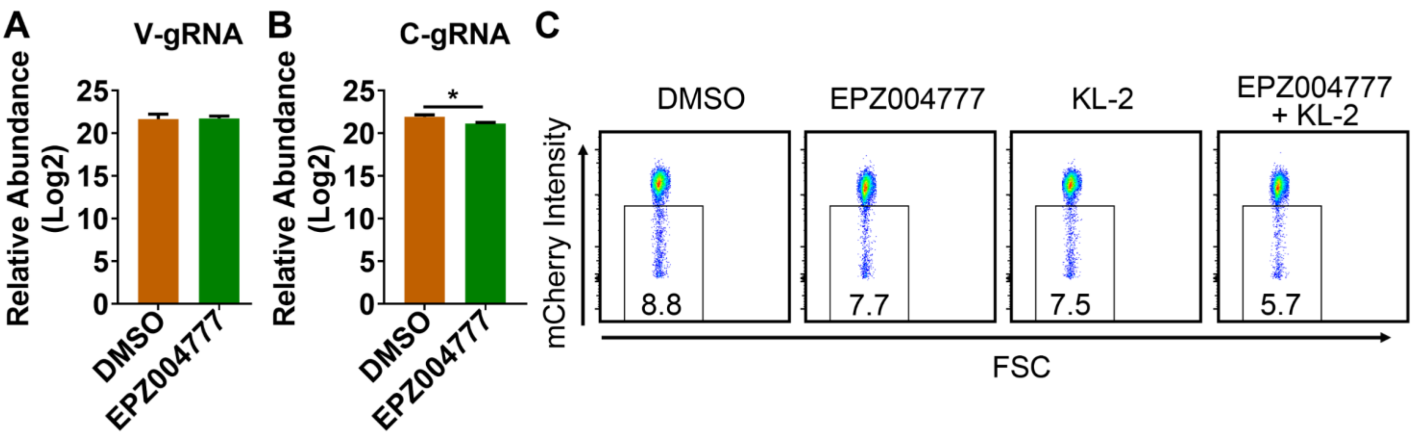

**Table S1 The relative abundance comparison of proteins identified between V and C regions that are known to play roles in SHM in B cells**

| Description | Symbol | V/C_FC_Log2 | V/C_Pvalue (-Log2) |
|---|---|---|---|
| Beta-catenin-like protein 1 | CTNNBL1 | 0.0 | 0.2 |
| FACT complex subunit SPT16 | SUPT16H | 0.0 | 0.3 |
| FACT complex subunit SSRP1 | SSRP1 | -0.1 | 1.2 |
| Germinal-center associated nuclear protein | MCM3AP | -0.5 | 1.9 |
| Polypyrimidine tract-binding protein 2 | PTBP2 | 0.0 | 0.0 |
| Polypyrimidine tract-binding protein 3 | PTBP3/ROD1 | -0.1 | 0.3 |
| Protein HIRA | HIRA | 1.1 | 4.1 |
| Single-stranded DNA cytosine deaminase | AICDA | 0.7 | 0.9 |
| Transcription elongation factor SPT4 | SUPT4H1 | -0.8 | 3.0 |
| Transcription elongation factor SPT5 | SUPT5H | 0.4 | 5.6 |
| Transcription elongation factor SPT6 | SUPT6H | -1.0 | 1.2 |

**Table S2 The information on the primers used for ChIP-qPCR in this study**

| Primer Name | | Sequence (5'-3') | Assay |
|---|---|---|---|
| TSS | Fw | GAGGACTCTGGGTTTGGTGA | ChIP q-PCR |
| | Rev | CTCACCCTGGGAGGTGTATG | |
| V region | Fw | AGCCTACAACGTCAACATCAAACTC | ChIP q-PCR |
| | Rev | GTTTCATAGATCTCCCGGAAGG | |
| up 500bp V | Fw | AATTCAGGGTCCAGCTCACA | ChIP q-PCR |
| | Rev | ATCCGCATTCCTGAGACACT | |
| Cμ | Fw | CCGATGTCTACTTGCTGCCACC | ChIP q-PCR |
| | Rev | GTCAGGATGCTGTGGGCGAAGT | |
| Cμ down 500bp | Fw | CTCAGTGGCTTCTAGAAACCCCTG | ChIP q-PCR |
| | Rev | CATGGGTTAGCACGTCTCTGT | |
| Sμ | Fw | CCATTTGAAGGAGAGGTCGC | ChIP q-PCR |
| | Rev | TGGGGCTTGGTATGTTCTCA | |

**Table S3 The information on the antibodies used in this study**

| Antibody | Company | Catalog | Assay |
|---|---|---|---|
| Anti-H3 | Abcam | ab1791 | ChIP, WB |
| Anti-H3K79me1 | Abcam | ab2886 | ChIP |
| Anti-H3K79me2/3 | Abcam | ab2621 | ChIP |
| Anti-biotin | Cell Signaling Technology | #7075 | WB |
| Anti-Flag | Millipore-Sigma | F1804 | ChIP, WB |
| Anti-ELOB | Bethyl | A304-008A | ChIP |
| Anti-CHD6 | Bethyl | A301-221A | ChIP |
| Anti-KAP1 | Bethyl | A300-274A | ChIP |
| Anti-ISY1 | Bethyl | A304-927A | ChIP |
| Anti-CWC15 | Bethyl | A304-900A | ChIP |
| Anti-HMGB2 | ABclonal | A17360 | ChIP |
| Anti-MacroH2A1 | Millipore-Sigma | ABE215 | ChIP |
| Anti-RBBP4 | ABclonal | A1490 | ChIP |
| Anti-RUVBL2 | ABclonal | A1905 | ChIP |
| Anti-MCM5 | ABclonal | A13514 | ChIP |
| Anti-MCM3 | ABclonal | A1060 | ChIP |
